# Multi-ancestry Transcriptome-Wide Association Study Reveals Shared and Population-Specific Genetic Effects in Alzheimer’s Disease

**DOI:** 10.1101/2025.11.03.686160

**Authors:** Xinyu Sun, Makaela Mews, Nicholas R. Wheeler, Penelope Benchek, Tianjie Gu, Lissette Gomez, Nicholas Ray, Christiane Reitz, Adam C. Naj, Jennifer Elizabeth Below, Giuseppe Tosto, Mario Cornejo-Olivas, Goldie S. Byrd, Briseida E. Feliciano-Astacio, Katrina Celis, Farid Rajabli, Brian W. Kunkle, Margaret A. Pericak-Vance, Jonathan L. Haines, Anthony J. Griswold, William S. Bush

## Abstract

Alzheimer’s disease (AD) risk differs across ancestral populations, yet most genetic studies have focused on non-Hispanic White (NHW) cohorts. We conducted a multi-population transcriptome-wide association study (TWAS) using whole-blood RNA-seq and genotype data from NHW (n=235), African American (AA; n=224), and Hispanic (HISP; n=292) MAGENTA participants. Using SuShiE for multi-population cis-eQTL fine-mapping, we identified credible sets for 8,748 genes, improving fine-mapping precision relative to analyses using fewer populations. cis-eQTL effects were largely shared across populations, with a subset showing population-specific regulation. We performed population-stratified TWAS of AD and inverse variance-weighted meta-analysis, followed by gene-level TWAS fine-mapping (MA-FOCUS), prioritizing nine genes (FDR<0.05, PIP>0.8), including established AD loci (*BIN1, PTK2B, DMPK*) with broadly consistent effects across populations. At *BIN1*, fine-mapped cis-eQTL variants used in the TWAS prediction model highlighted rs11682128, which is only modestly correlated with the GWAS index SNP rs6733839 (*r*^2^ ≈ 0.34), demonstrating how integrating eQTL fine-mapping with TWAS can refine signals beyond sentinel GWAS variants. We also identified an association between *COG4* expression and AD in NHW, implicating Golgi-related pathways. Using independent SuShiE-derived models from TOPMed MESA (PBMC), several signals replicated directionally across ancestries, with the strongest statistical support in NHW. Overall, multi-population eQTL fine-mapping improves model interpretability and helps resolve shared and population-specific regulatory mechanisms relevant to AD.

## Introduction

Alzheimer’s Disease (AD) is the most common form of dementia and affects millions worldwide, with its prevalence expected to increase significantly as global populations age^1^. AD is heritable, though AD risk varies substantially across ancestral populations, with African Americans having approximately double the risk and Hispanics having about one and one-half times the risk compared to non-Hispanic Whites^1,2^. Recent and ongoing ascertainment efforts have increased participation of African American and Hispanic individuals in genetic studies, which have led to multiple ancestry-specific GWAS^3^, and a recent multi-ancestry GWAS meta-analysis^4^. These studies (along with detailed analyses of APOE-ε4 effects^5^) have revealed substantial heterogeneity in effect sizes of established AD risk variants^4^, and different genetic architectures in AD risk across ancestries.

Growing recognition that AD involves coordinated dysregulation of both peripheral and central immune systems has shifted the field toward viewing AD as a systemic disorder. As highlighted by Bettcher et al.^6^, mounting clinical and mechanistic evidence reveals dynamic crosstalk between the peripheral immune system and the central nervous system, suggesting that immune perturbations in blood may not only reflect, but also influence, AD pathogenesis. This systems-level perspective underscores the value of studying whole blood as an informative, accessible tissue for understanding disease mechanisms across diverse populations.

Multiple transcriptome-wide association studies (TWAS) of AD have been conducted to extend GWAS results to functional effects in brain and blood^7–17^. In principle, TWAS jointly models the effects of multiple (typically non-coding) variants on genetically predicted gene expression and tests its association with a phenotype, providing gene-level, biologically interpretable hypotheses that can complement single-variant GWAS findings. Various multi-variable modeling strategies (such as Elastic net^18^ and LASSO^19^) have demonstrated gene expression is predominantly driven by a sparse set of genetic variants^20–22^. Recent methodological advances in Bayesian fine-mapping^23,24^ have also led to approaches that use fine-mapped cis-eQTLs as variables for gene expression prediction^21,25^. Bayesian fine-mapping methods such as SuSIE^23,24^ quantify uncertainty by defining credible sets of highly correlated variants, which enables more robust interpretation in the presence of complex linkage disequilibrium (LD) structures^23^. Notably, all of these approaches require population-specific eQTL reference panels^26^ that have distinct eQTL effects specific to a particular population and also match the LD structure of the target GWAS dataset. Due to the scarcity of non-European eQTL datasets, most TWAS have been conducted in non-Hispanicwhite populations only.

Several multi-ancestry TWAS methods have been developed to leverage expression data across populations. METRO employs a joint likelihood-based inference framework that uses marginal eQTL estimates trained independently within each ancestry and then combines them in a joint likelihood framework to estimate ancestry-specific contributions while explicitly modeling uncertainty in the input estimates.^27^, and has been applied to identify genes associated with Alzheimer’s disease and cognitive function using European and African ancestry expression data^28^. MATS uses a random effects model to jointly analyze samples from multiple populations, distinguishing between shared, ancestry-specific, and subject-specific expression-trait associations, but relies on separately constructed prediction models^29^. TESLA integrates an eQTL dataset with multi-ancestry GWAS using an optimal linear combination of association statistics to exploit shared effects while accommodating heterogeneity^30^. A common feature of these approaches is that expression prediction models or eQTL data are trained independently within each ancestry before being combined in the association testing framework.

In contrast, SuShiE (Sum of Shared Single Effects) was recently developed for multi-ancestry fine-mapping of molecular QTLs, leveraging LD heterogeneity across populations to improve fine-mapping precision while simultaneously inferring cross-ancestry effect size correlations and providing prediction weights for each population^31^. In this work, we perform multi-ancestry fine-mapping of whole-blood eQTL effects and conduct a multi-ancestry TWAS of AD within reported African American (AA), non-Hispanic White (NHW), and Hispanic (HISP) participants in the MAGENTA study^32^. We first build population-specific gene expression prediction models, then perform population-stratified TWAS, integrate these associations via sample-size-weighted meta-analysis, and fine-map putative causal genes. Finally, we compare dense (all cis-SNPs) versus sparse (credible set-based) models.

## Material and Methods

### eQTL Reference Panel Study Participants

The *Multi-Ancestry Genomics, Epigenomics, and Transcriptomics of Alzheimer’s* (MAGENTA) study accessed existing sample collections that were part of multiple previous NIA-funded research projects^32^. Some legacy samples were drawn from studies that ascertained White or African American participants. All other samples were drawn from more recent studies including the Puerto Rican Alzheimer’s Disease Initiative, the Cuban American Alzheimer’s Disease Initiative, the Peru Alzheimer’s Disease Initiative, and the Research in African-American Alzheimer’s Disease Initiative. All participants and/or their consenting proxy provided written informed consent as part of the study protocols approved by the site-specific Institutional Review Boards.

### Population Descriptors and Study Group Definitions

Consistent with recent recommendations^33^, we define our rationale and definition for the population descriptors in this study. MAGENTA participants were recruited across multiple studies that differed in catchment geography (U.S., Peru), terms used for study recruitment and outreach, and cultural context. As a result, demographic fields and participant descriptors varied across individuals.

To enable cross-group comparisons while acknowledging differences in genetic ancestry and admixture, we used a combination of geographic and self-identity terms as follows:

- “Non-Hispanic White” (NHW): individuals ascertained in North Carolina, Tennessee, and South Florida where participants self-identified as non-Hispanic White or were categorized as not Hispanic in legacy datasets where recontact is impossible.
- “African American” (AA): individuals ascertained in Northeast Ohio, North Carolina, and South Florida using descriptors intended to reach self-identified Black/African American participants.
- “Caribbean Hispanic”: individuals ascertained in the U.S. using descriptors intended to reach individuals with Cuban or Puerto Rican heritage, reflecting similar admixture patterns and environmental exposures.
- “Peruvian Hispanic”: individuals ascertained in Lima, Peru and surrounding areas.

For consistency with prior published GWAS studies^4,34,35^, we combined the Caribbean Hispanic and Peruvian Hispanic into a single “Hispanic” group to facilitate our TWAS analyses. These labels reflect recruitment context and self-identification, and are 91% concordant with ancestry-based clusters; downstream genetic analyses incorporate ancestry-informative covariates to account for population structure.

### Genotyping and Quality Control

All samples were genotyped using either the Illumina MEGA v2 or v3 multi-population array and subsequently imputed using TOPMed reference panels v2 (build 38) corresponding with self-identified ancestry and/or ascertainment criteria. We restricted our analysis to variants directly genotyped by the array and variants with an imputation quality control score of *R*^2^ > 0.8. Genotyping data was processed using the ADSP xqtl-protocol release 0.1.1. All analyses were performed stratified by group yielding a total of three analyses. Relatedness was assessed using kinship analysis generated by the KING software. Subsequently, we removed samples to eliminate relatedness (kinship coefficient ≥ 0.0625) and maintained samples by prioritizing inclusion of cognitively normal individuals with advanced age, diagnosed AD cases, and control samples with rarer APOE genotypes (ε3/4 or ε4/4). We concluded our genotyping quality control using PLINK2.0 to filter variants with minor allele counts (MAC) of 10, Hardy-Weinberg equilibrium of 1* 10 ^− 8^, variant-level missingness of 0.1, and sample-level missingness of 0.1.

Post-QC, 224 AA, 235 NHW, 292 HISP samples from MAGENTA, with both whole-blood expression and genotype data, were analyzed. The gene expression was measured in whole blood using RNA-seq. We included the common protein-coding genes that were available in all three populations. In total, we retained 14,436 genes. Major Histocompatibility Complex (MHC) region from chromosome 6p (∼24-34 Mbp) was excluded from the study, due to the complex LD structure. In the end, we analyzed 14,211 protein-coding genes.

### RNA Processing

Sample preparation and RNA sequencing of whole blood samples have been previously described^32^. Briefly, RNA was extracted from previously frozen whole blood samples, quality controlled with RNA integrity (RIN) scores >= 5, prepared using the NuGEN Universal Plus mRNA-Seq with globin and ribosomal depletion, and sequenced using 125bp paired end Illumina HiSeq3000. RNA sequencing FASTQ files were processed using the STAR aligner (v.2.7.10a), quality controlled using Picard, and quantified using RSEM (v.1.3.3) using the following reference files:

GRCh38_full_analysis_set_plus_decoy_hla.noALT_noHLA_noDecoy_ERCC.fast a, Homo_sapiens.GRCh38.103.chr.reformatted.collapse_only.gene.ERCC.gtf, and hg38_GENCODE.v38.bed (see ADSP xqtl-protocol 0.1.1 for full details).

Genes with a TPM expression level of 10% or less in over 20% of samples and/or genes with read counts less than 6 among more than 20% of samples were excluded from the analysis. In addition, samples detected as outliers using relative log expression (RLE), hierarchical clustering, and D-statistics were removed. Finally, TPM data was normalized using the Trimmed Mean of M-value (TMM) method.

### Bulk RNA-seq Deconvolution and Hidden Factor Detection

As we used bulk whole blood RNA data for our analysis, we performed CIBERSORTx deconvolution using the European-based LM22 reference panel to estimate relative cell-type proportions in each sample^36^. We performed this analysis using all samples combined. We included estimated cell-type proportions for 14 estimated cell types as covariates in our analysis based on an identified significant difference in relative cell-type proportions within each population or across AD status (Kruskal-Wallis Test p < 0.05). The 14 estimated cell-type proportions are Neutrophils, CD4+ T cells-Memory resting, B cells - Naïve, B cells - Memory, Macrophages - M1, Mast cells - activated, T cells - follicular helper, Plasma cells, T cells - CD4 naive, Regulatory T cells - Tregs, Macrophages - M0, NK cells - activated, Monocytes, and NK cells - resting.

We also performed Hidden Factor Analysis using the Marchenko-Pastur PCA approach as described in the ADSP xqtl-protocol 0.1.1. The approach applies the Marchenko-Pastur distribution to determine the optimal number of principal components to retain by modeling the eigenvalues of random covariance matrices. Components with eigenvalues above the Marchenko-Pastur threshold are considered to represent meaningful biological variation, while those below it are attributed to random noise. This technique ensures that only components capturing substantive biological variation are retained for downstream analyses, avoiding overfitting while enhancing the interpretability of confounder correction in our QTL mapping. In total, there are 42, 41 and 49 hidden factor PCs used in NHW, AA and HISP population respectively.

### AD GWAS Summary Statistics

We accessed the publicly available clinically diagnosed AD GWAS summary statistics for each population. All summary statistics were derived from models adjusted for age, sex, and population substructure principal components. We lifted coordinates from GRCh37/hg19 using UCSC `liftOver`^37^ as needed.

We accessed summary statistics from the Ray et al. African American GWAS, which included 2,784 AD cases and 5,222 controls (total n=9,168)^34^. We also accessed summary statistics from the Hispanic subset of Rajabli et al., comprising 3,005 AD cases and 5,894 controls (total n=8,899)^4^. For the non-Hispanic White population, we used Stage I data from Kunkle et al., which included 21,982 AD cases and 44,944 controls (total n=63,926)^35^.

### Harmonization of GWAS and Expression Prediction Reference

To ensure consistency and improve the integration of GWAS summary statistics with our gene expression prediction reference panel, we performed several harmonization procedures. First, we ensured the reference and alternative alleles of each variant were defined consistently across the GWAS and the reference panel. If the reference and alternative alleles were swapped between the GWAS and the reference panel, we inverted the GWAS Z-score so that its sign matched the alternative allele in the reference panel. We also identified and corrected strand flips, where alleles were reported on opposite DNA strands between datasets, by checking for complementary base pair substitutions and adjusting allele coding accordingly.

Second, we performed summary statistics imputation to maximize SNP coverage between the GWAS and prediction models. Publicly available GWAS vary in sequencing platforms, imputation panels, and genome builds, resulting in inconsistent variant availability across studies^21^. Since prediction model performance depends on the availability of constituent variants in the GWAS, missing variants can impair the ability to properly capture expression variation patterns. We followed the PredictDB GTEx v8 GWAS imputation pipeline to impute missing variant Z-scores according to the MAGENTA genotype reference panel for each population. This pipeline also performs strand flip quality control as part of its harmonization procedures. A total of 1,252,104, 1,253,727, and 1,081,547 variants were imputed for AA, EUR, and HISP GWAS respectively, representing 19.07% of total variants on average.

Third, only unambiguous SNPs-those that do not involve substitutions of complementary base pairs (i.e., excluding A/T and C/G variants)-were retained to avoid potential strand ambiguity.

### Statistical methods

#### SuShiE

We applied SuShiE^31^ to perform multi-ancestry cis-eQTL fine-mapping using individual-level gene expression and genotype data from all three populations. SuShiE jointly models data across ancestries to infer credible sets of putative causal variants and estimate population-specific cis-eQTL effect sizes (weights) for predicting genetically regulated gene expression (GReX).

Following empirical convention^23,24,31^, we set the maximum number of credible sets to *L*= 10 for each protein-coding gene and specified a posterior inclusion probability (PIP) threshold of 0.9 for SNPs to be included in credible sets; default settings were used for all other parameters (Table S3) except that we did not apply an additional minor allele frequency (MAF) filter because variants had already been filtered to *MAC* ≥ 10 during genotype QC. Each credible set is designed to capture a homogeneous and distinct causal signal; to ensure this, we applied a purity filter removing credible sets with sample-size-weighted minimum absolute pairwise correlation below 0.5.

SuShiE produces population-specific cis-eQTL weights, where SNPs with large-magnitude weights correspond to credible set variants. This variable selection approach reduces TWAS estimation errors by mitigating the influence of weak prediction instruments^38^. SuShiE also estimated cis-SNP heritability for each gene within each population using the limix Python package.

Cis-eQTL weighted prediction modeling was performed for each protein-coding gene using SNPs within a ± 500kb region (500kb before TSS, and 500kb after TES), adjusting for age, (self-reported & QC on genetic data) sex, top three genetic PCs, cell-type proportions and hidden factors.

While the posterior inclusion probability (PIP) indicates the probability that a SNP is causal in the joint multi-ancestry model, it does not directly distinguish whether the underlying effect is population-specific or shared with heterogeneous directions across ancestries. To address this, SuShiE also estimates a prior effect correlation parameter (ρ) for each single effect, representing the inferred co-variation of causal effect sizes across ancestries within each gene. To further characterize genes exhibiting strong negative correlations (ρ < -0.5) in the primary single effect (CS1), we classified effect patterns based on the posterior SNP weights. For each ancestry pair, we computed the sum of posterior weights across CS1 variants and classified genes as exhibiting “opposing direction” effects if both ancestries showed meaningful effects (|effect| ≥ 0.05) of opposite sign, or “population-specific” effects if one ancestry showed a meaningful effect while the other was near-zero (|effect| < 0.05). The near-zero threshold of 0.05 was chosen conservatively, below the 25% percentile of absolute summed weight (∼0.0582), to ensure population-specific classification requires a truly negligible effect in one ancestry.

#### FUSION

FUSION estimates the association between genetically regulated gene expression (GReX) and Alzheimer’s disease using GWAS summary statistics. The TWAS test statistic (*Z*_*TWAS*_) is computed as a weighted combination of GWAS SNP-level Z-scores (*Z*_*gwas*_), where the weights (*W*_*snp*_) are derived from the SNPs’ effects on gene expression. To account for correlation structure among SNPs, the test statistic is normalized by the square root of the weighted linkage disequilibrium (LD) matrix:

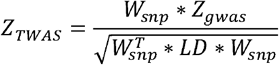

We followed the FUSION^39^ pipeline and used MAGENTA genotype data as the LD reference panel for each population-stratified TWAS analysis. SuShiE-derived population-specific cis-eQTL effect sizes were used as prediction weights to estimate GReX for each gene.

We constructed two types of prediction models from SuShiE output. Dense models used weights for all cis-SNPs, which SuShiE computes for every gene regardless of whether credible sets are identified. Sparse models used weights restricted to only SNPs within credible sets that passed purity filtering (n=8,748); these were available only for genes with at least one credible set. In our main TWAS analyses, we used dense models to maximize the number of testable genes (14,127 models successfully trained). Sensitivity analyses using sparse models for genes with credible sets are reported separately to assess the impact of variable selection on TWAS results.

There are 14,127, 14,116, and 13,742 gene models successfully tested for NHW, AA, and HISP respectively. There are 13,731 common models tested across all populations. We set the FDR’s q-value threshold to 0.05 (*q*_*c*_ < 0.05) to identify AD-related gene candidates.

#### Meta-analysis of population-stratified TWAS associations

As suggested by Bhattacharya et al.^26^, inverse-variance weighting (IVW) meta-analysis was performed on three population-stratified TWAS associations to share TWAS associations across all populations. To estimate the TWAS effect size in FUSION code, we adopted the formulation from S-PrediXcan^40^.

#### TWAS Effect Size Estimation

FUSION^41^ natively outputs Z-scores for gene-trait associations but does not provide effect sizes. To enable inverse-variance weighted meta-analysis, we extended FUSION to compute TWAS effect sizes (*β*_*TWAS*_) and standard errors following the S-PrediXcan framework^40^. For standardized genotypes, the TWAS effect size is computed as:

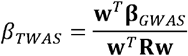

where **w** is the vector of cis-SNP prediction weights, **β**_*GWAS*_ is the vector of GWAS effect sizes, and **R** is the LD correlation matrix from the reference panel. This formulation is mathematically equivalent to S-PrediXcan when using standardized genotypes, as FUSION’s correlation matrix corresponds to S-PrediXcan’s covariance matrix scaled by unit variance. The standard error was derived from the Wald statistic relationship: *SE* = *β*_*TWAS*_/*Z*_*TWAS*_.

We performed an IVW meta-analysis across ancestry-stratified TWAS results and controlled for multiple testing using FDR. Genes with (*q*_*c*_ < 0.05) were considered significant in the meta-analysis. We note that ancestry-pairwise concordance of TWAS associations was modest, consistent with prior work^26^, which motivates caution in interpreting meta-analyzed signals as universally shared.

### TWAS Fine-mapping

TWAS may nominate multiple genes, but some of these genes could be non-causal due to sharing of eQTLs or gene co-regulation. We applied MA-FOCUS^42^ to prioritize putative causal genes by correcting for correlation structures induced by linkage disequilibrium (LD) and prediction weights.

MA-FOCUS is a multi-ancestry extension of FOCUS that takes population-specific triplets of inputs-LD reference panels, cis-eQTL prediction weights, and GWAS summary statistics-to perform joint fine-mapping across ancestries. For each LD block, MA-FOCUS internally computes TWAS statistics and performs fine-mapping to produce both population-specific posterior inclusion probabilities (*PIP*_*pop*_) and a multi-population posterior inclusion probability (*PIP*_*ME*_). The *PIP*_*ME*_ represents the probability that a given gene is causal for the trait, computed jointly across all population-specific datasets under the assumption that causal genes are shared across ancestries^42^.

Population-specific genotype data from NHW, AA, and HISP groups in the MAGENTA study served as LD reference panels. We performed fine-mapping on LD blocks containing at least one significant gene (*q*_*c*_ < 0.05) using default parameters. For each LD block, MA-FOCUS calculated PIPs and derived 80% credible sets (ρ = 0.8) of putative causal genes. To prioritize genes with the strongest evidence of causality, we classified a significant TWAS association as a high-confidence causal gene if it achieved either a population-specific *PIP*_*pop*_ > 0.8 in its corresponding population-stratified analysis, or a multi-population *PIP*_*ME*_ > 0.8 in the IVW cross-ancestry meta-analysis.

While PIPs quantify the probability that a gene is causal, they do not directly indicate whether the underlying effect is population-specific (non-zero in one ancestry but zero in others) or shared with heterogeneous directions across ancestries. To distinguish these scenarios, we examined the population-specific TWAS Z-scores and *PIP*_*pop*_ values provided by MA-FOCUS. Genes with high *PIP*_*ME*_ but discordant signs in population-specific TWAS Z-scores suggest shared causal genes with opposite effect directions.

### Independent Multi-ancestry Expression Reference Panel

In our analyses, we re-evaluated the significant TWAS associations from our discovery dataset (MAGENTA) using the publicly released SuShiE-derived weights from the TOPMed MESA visit-1 mRNA dataset^31,43,44^. As we did not have access to the individual level genotype data of the MESA samples, we used high-coverage 1000 Genomes data to generate a LD reference for each ancestry^45^. The MESA dataset contained expression measurements from 956 participants (402 European American, 175 African American, and 277 Hispanic American individuals) with corresponding whole genome sequencing data. The mRNA expression was quantified in peripheral blood mononuclear cells (PBMCs), covering 21,747 protein-coding genes. The original analysis adjusted for 15 gene expression principal components, 10 genotype principal components, age, sex, and assay lab as covariates. These SuShiE-derived prediction models (version 1) were accessed at https://zenodo.org/records/10963034.

### Functional Annotation Using FILER

To characterize the regulatory potential of fine-mapped eQTL variants in additional candidate genes (*COG4*) that has not been reported, we queried the Functional genomics repository (FILER) database^46^, which integrates functional genomic annotations from multiple consortia including ENCODE, Roadmap Epigenomics, and EpiMap.

#### Chromatin State Annotation

Chromatin states were retrieved via FILER from the EpiMap, which provides ChromHMM-based chromatin state predictions across 833 biosamples^47^. The 18-state ChromHMM model classifies genomic regions into functional states. We focused on enhancer states including active enhancers (EnhA1, EnhA2), genic enhancers (EnhG1, EnhG2), and weak enhancers (EnhWk) in blood and immune cell types.

#### Chromatin Interaction Analysis

As supplementary functional evidence, we queried promoter-capture Hi-C (pcHi-C) data from the Javierre et al. study^48^, which profiled chromatin interactions between promoters and distal elements across 17 human primary hematopoietic cell types. Significant interactions were defined using the CHiCAGO algorithm with a score threshold of 5^49^.

## Results

### MAGENTA Study Overview

We used the MAGENTA dataset as the eQTL reference, comprised of African American (AA, n=224), non-Hispanic White (NHW, n=235), and a combined Hispanic group (HISP, n=292). The Hispanic group contains Caribbean and Peruvian sub-cohorts to align with available AD GWAS summary statistics. Case/control status was approximately balanced across groups (∼50% each), and age distributions were comparable (means ∼76.6-77.5 years), supporting cross-population comparability for downstream fine-mapping and TWAS analyses (Table S1).

We found that cis-eQTL effects were largely consistent whether or not Alzheimer’s disease (AD) status was included as a covariate in the MAGENTA datasets (Figure S3, Table S5, Figure S5), consistent with previous studies^50–52^.

### From multi-ancestry fine-mapping to interpretable TWAS: workflow overview

Our multi-population workflow (Figure 1) integrates MAGENTA expression and genotype data with population-matched AD GWAS to generate interpretable TWAS associations. SuShiE multi-ancestry fine-mapping yields population-specific GReX models that we test with FUSION within each ancestry and meta-analyzed across populations, followed by MA-FOCUS gene-level fine-mapping of TWAS hits to identify putative causal genes. A detailed dense versus sparse comparison demonstrating that credible-set variants recapitulate dense-model TWAS signals is presented in the “Sparse Models Using Fine-mapped eQTLs” subsection. Overall, we showed this workflow increases eQTL resolution and enables variant-resolved interpretation of of the expression model instruments underlying TWAS.

**Figure 1:**
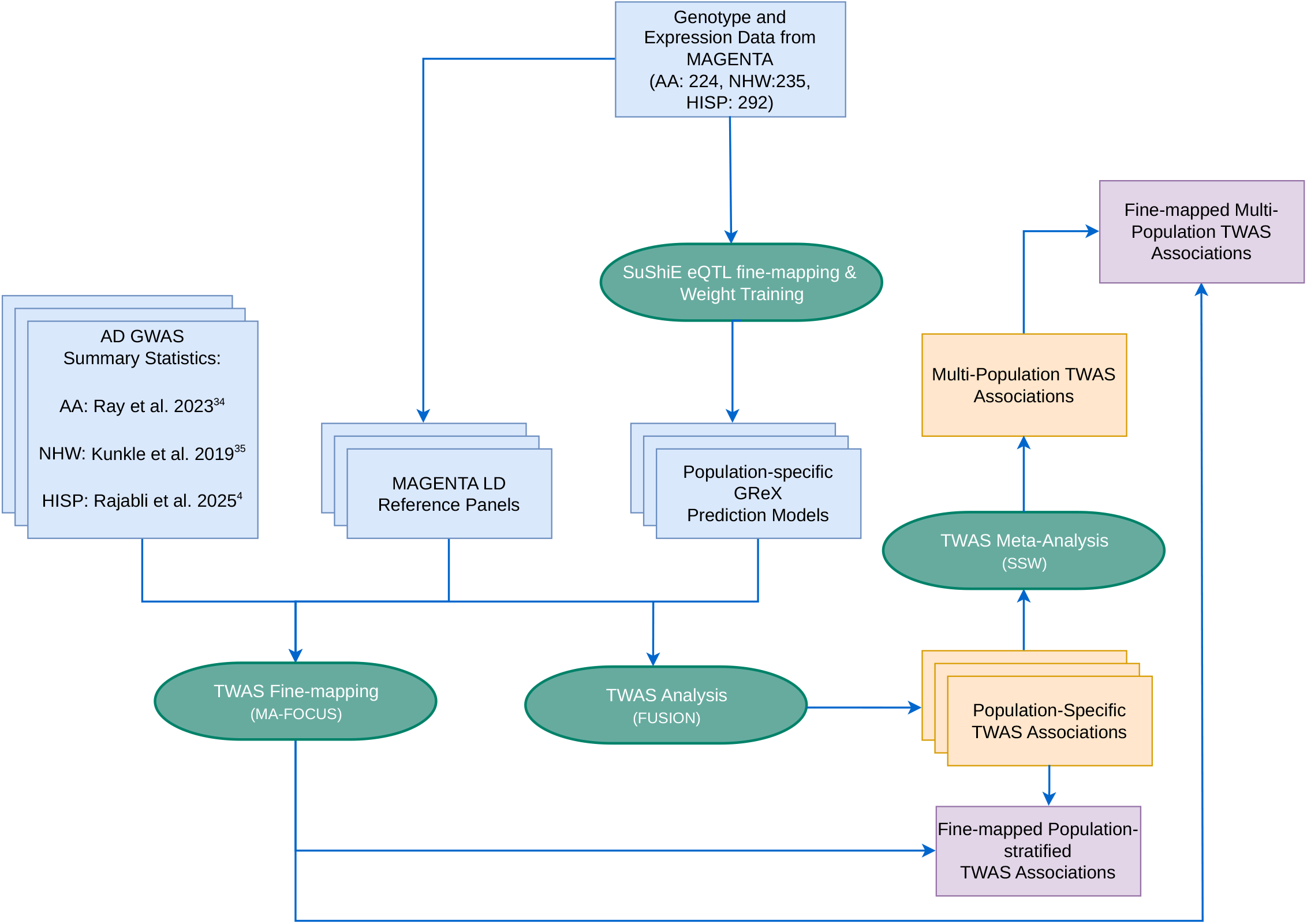
Multi-population TWAS analytical workflow for Alzheimer’s Disease. The workflow integrates MAGENTA genotype and expression data (n=751; AA: 224, NHW: 235, HISP: 292) with published AD GWAS summary statistics across three population groups: Hispanic (Rajabli et al. 2025^4^), African American (Ray et al. 2022^34^), and non-Hispanic White (Kunkle et al. 2019^35^). SuShiE is applied for multi-population eQTL fine-mapping to create population-specific GReX prediction models, which are then used in FUSION analysis alongside GWAS summary statistics and population-matched reference panels. Population-stratified TWAS results are combined through Inverse-Variance Weighted (IVW) meta-analysis to produce multi-population TWAS associations. Fine-mapping of these results using MA-FOCUS identifies the most likely causal genes.

### Multi-population fine-mapping improves credible set precision and identifies shared eQTL effects

Using MAGENTA whole-blood RNA-seq and genotypes from AA, NHW, and HISP (14,211 shared protein-coding genes), we applied SuShiE multi-ancestry fine-mapping and identified credible sets (CS) for 8,748 genes. The mean number of CS per gene (with at least one CS) was 1.95 (median 2), with very few genes requiring more than seven CSs, supporting our choice of maximum number of expected CS, *L* = 10. Credible sets were compact (median 4 SNPs; Figure S1), consistent with a sparse cis-regulatory architecture^20–22,53,54^.

Including all three populations improved fine-mapping precision relative to any two-population analysis (NHW-AA, NHW-HISP, and AA-HISP) (Table 1); median SNPs per CS decreased from 6 to 4 (-33%), consistent with reduced LD-induced ambiguity across ancestries, the mean number of CS per gene with at least one CS increased from 1.36 to 1.95 (+43%), and the fraction of SNPs with *PIP*> 0.9 rose from 0.018 to 0.024 (+32%), indicating improved confidence in variant-level fine-mapping. These findings are consistent with the benefits of multi-ancestry fine-mapping results reported by the PAGE consortium^55^.

**Table 1:** Improved fine-mapping precision of shared cis-eQTLs through multi-population analysis.

| Metric | Average of Any Two Populations | Three Populations (NHW+AA+HISP) | Improvement |
| --- | --- | --- | --- |
| Median SNPs per credible set | 6 | 4 | 33% reduction |
| Mean credible sets per gene | 1.36 | 1.95 | 43% increase |
| Percent of eQTL had PIP > 0.9 | 0.018 | 0.024 | 32% increase |

<sup>a</sup>This table demonstrates the benefits of incorporating three diverse populations (non-Hispanic White, African American, and Hispanic) in SuShiE analysis compared to using only two populations. The tri-population approach resulted in more precise credible sets (33% fewer SNPs per set), identified more credible sets per gene with at least one CS (43% increase), and increased the proportion of SNPs with high posterior inclusion probability (32% increase in SNPs with $PIP > 0.9$ ). The denominator for “Percent of eQTL had PIP > 0.9” is the number of SNPs in genes with at least one credible set (NHW-HISP: 111,507; HISP-AA: 71,167; NHW-AA: 84,711; all three populations: 170,059). These findings support the sparse genetic architecture of gene expression regulation and highlight the value of diverse population inclusion in fine-mapping studies.

Following the approach of Lu et al.^31^, we evaluated cross-ancestry consistency of effects for the first SuShiE-reported single shared effect (CS1), which represented the largest single effect on gene expression. The prior effect correlation parameter (ρ)-which captures the model’s inferred co-variation of causal effect sizes across ancestries within each gene-was highly consistent for roughly 70% of genes, indicating broadly shared regulation (Figure 2). A subset of genes (∼30%) showed anticorrelation (ρ < -0.5), with 2,509 (AA-NHW), 2,990 (AA-HISP), and 2,954 (NHW-HISP) genes, and substantial pairwise overlap (1,223-1,708 genes) where effect sizes of SNPs on gene expression exhibited discordant effect directions across ancestry pairs. To characterize whether these negative correlations reflected opposing direction effects (substantial effects of opposite sign in both ancestries) or population-specific effects (substantial effect in one ancestry, near-zero in the other), we examined the posterior SNP weights for these genes. Across all population pairs, opposing direction effects predominated (AA-NHW: 67%, n=1,671; AA-HISP: 58%, n=1,732; NHW-HISP: 58%, n=1,721), with population-specific effects-where one ancestry has a meaningful effect while the other is near-zero-comprising 31-38% of negative-correlation genes.

**Figure 2:**
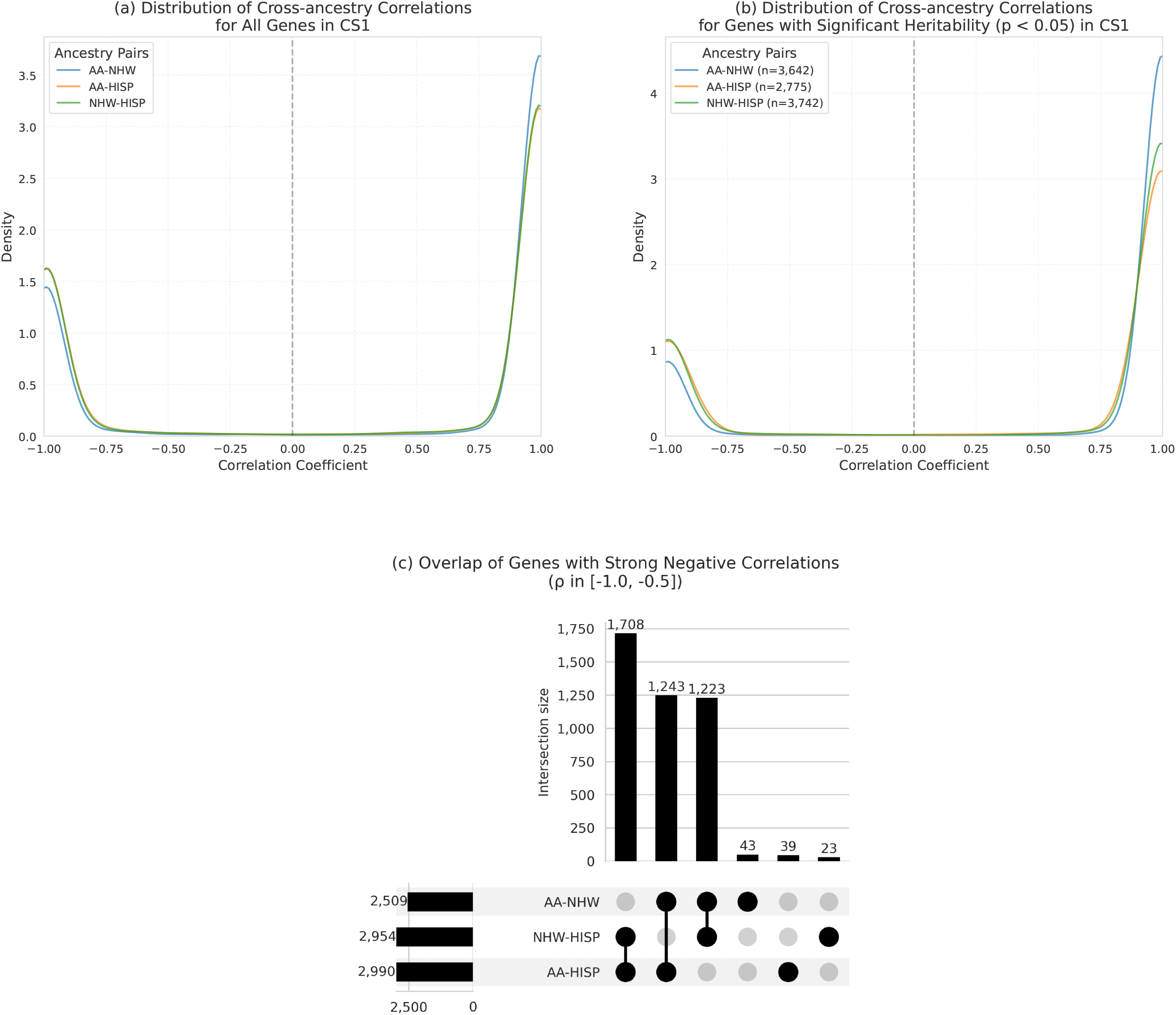
Cross-population analysis of eQTL effects in the first credible set (CS1). (a) Density plots showing the distribution of prior effect size correlations across population pairs (NHW-AA, NHW-HISP, and AA-HISP) for all genes (n=8,748) and (b) genes with significant heritability (p < 0.05). The bimodal distributions indicate predominantly consistent effects across populations (∼70% positive correlations) with some population-specific patterns. (c) UpSet Plot showing overlap of genes with strong negative correlations (r < -0.5) between population pairs. The substantial pairwise overlaps (1,223-1,708 genes) suggest that genetic effects typically differ in one population while remaining consistent between the other two, with HISP showing the most distinct patterns (1,708 overlapping genes with other pairs).

We then assessed the aggregate concordance of posterior eQTL weights for variants in CS1 across populations. Unlike the per-gene prior correlation, this measure summarizes the overall consistency of predicted SNP effect sizes-which represent the sum of contributions across all inferred single effects-across all genes. For all genes, correlations were moderate (AA-NHW r=0.526; AA-HISP r=0.379; NHW-HISP r=0.441), and increased when restricting to genes with significant cis-heritability in at least one population (AA-NHW r=0.689; AA-HISP r=0.484; NHW-HISP r=0.540) (Figure S2). These results support substantial sharing of cis-regulatory mechanisms across ancestries, with clearer concordance in genes under stronger genetic control, though notable population-specific effects remain.

### Population-stratified TWAS reveals both shared and group-specific AD gene associations

We applied MAGENTA-derived, population-specific weights in FUSION to AD GWAS from Kunkle et al. (NHW)^35^, Ray et al. (AA)^34^, and Rajabli et al. (HISP)^4^. We identified minimal inflations in our TWAS study, well within the acceptable limits (*λ*_*GC*_: NHW = 1.080, AA = 0.909, HISP = 0.979). At *q*_*c*_ < 0.05 (NHW: *P*< 1.4 * 10 ^− 4^ ; AA: *P* < 2.8 * 10 ^−6^ ; HISP: *P*< 3.8 * 10 ^−6^), we identified 40, 4, and 3 significant genes, respectively (Figure 3 (a-c); Table S10). Using MA-FOCUS to prioritize putative causal genes among the significant hits (*q*_*c*_ < 0.05), we identified high-confidence genes with *PIP* > 0.8: 12 in NHW and 2 each in AA and HISP.

**Figure 3:**
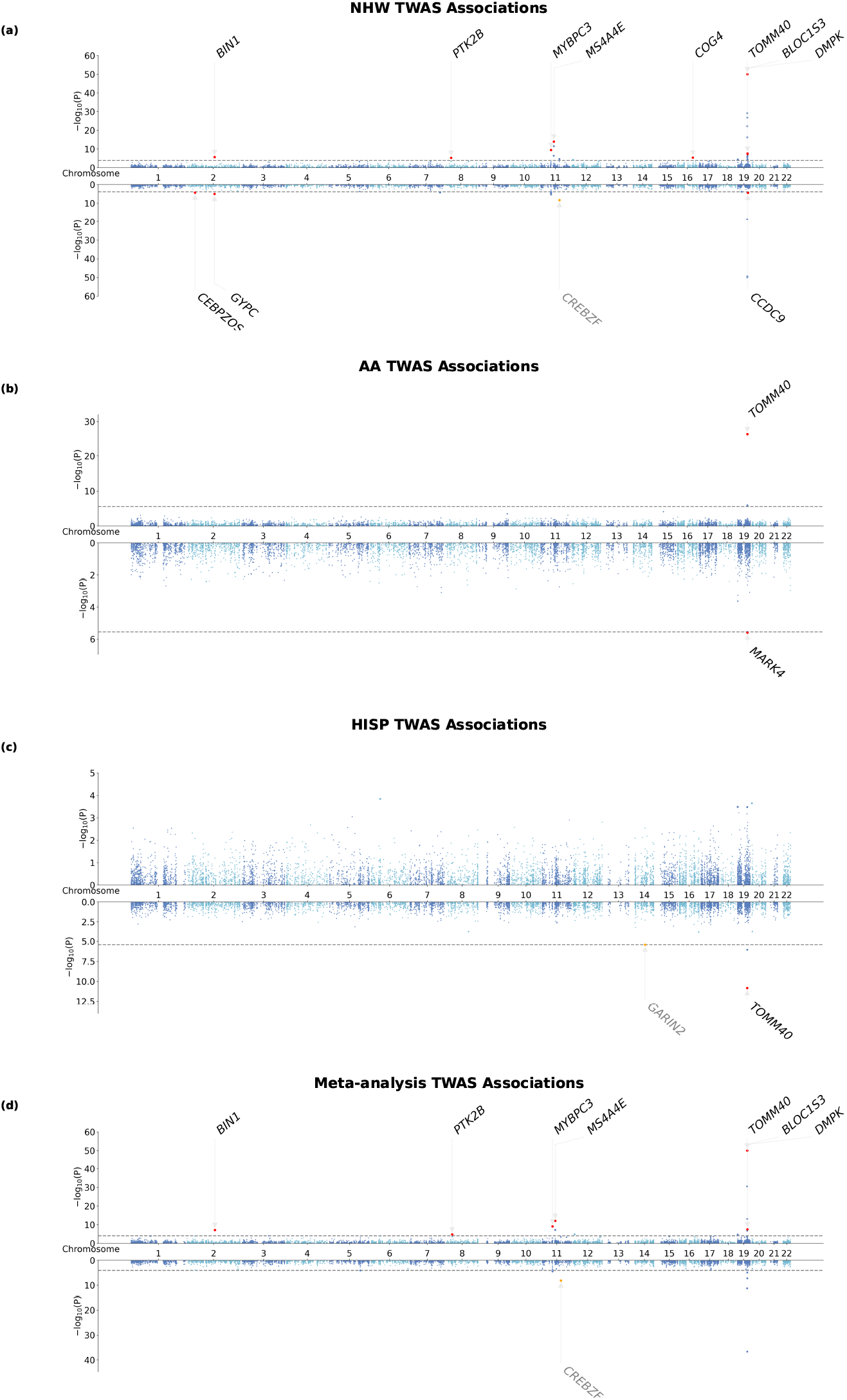
Population-stratified and meta-analyzed TWAS associations for Alzheimer’s Disease across diverse populations. The figure displays TWAS associations as Manhattan plots with -log10(P-value) on the Y-axis, with positive associations shown in the top panel and negative associations in the bottom panel. The gray dashed lines indicate the population-stratified or cross-ancestry meta-analyzed FDR threshold of 0.05, corresponding to *P*_*NHW*_ < 1.4 * 10 ^−4^ ; *P*_*AA*_ < 2.8 * 10 ^−6^ ; *P*_*HISP*_ < 3.8 * 10^−6^ ; *P*_*META*_ < 1.4 * 10^−6^ for each respective population. Red dots highlight genes that meet both statistical significance (*FDR* < 0.05) and fine-mapping criteria (*PIP*> 0.8), with gene symbols labeled for these prioritized associations. Each panel represents different population-stratified or meta-analyzed associations. Genes highlighted in orange dots and grey in gene symbols (*GARIN2, CREBZF*) reached TWAS significance but lack both significant prediction R^2^ in any population and identifiable eQTL credible sets from fine-mapping, suggesting these signals likely reflect horizontal pleiotropy. a) A Miami plot of NHW population-stratified TWAS associations; b) A Miami plot of AA population-stratified TWAS associations; c) A Miami plot of HISP population-stratified TWAS associations; d) A Miami plot of IVW meta-analyzed TWAS associations

We first highlight within-population signals (*q*_*c*_ < 0.05, population-specific *PIP*> 0.8) that did not reach significance in the multi-population meta-analysis. In NHW, two genes outside known AD loci were significant: *CEBPZOS* and *COG4* (Figure 3 (a)). *CEBPZOS* showed a negative association (*Z*_*NHW*_ = -4.076, *P*= 4.58 * 10 ^−5^). SuShiE fine-mapping identified 11 SNPs in 5 credible sets; CS1-CS2 exhibited near-perfect cross-population effect concordance 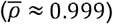, whereas CS3-CS5 were population-specific (anticorrelated). In particular, NHW’s effect size correlation from CS4-5 exhibited strong anticorrelation than HISP and AA (Figure S7). Expression was strongly predictable in NHW (*R*^2^ = 0.225) and modestly but significantly in AA (*R*^2^ = 0.0246, *P* = 0.0106). Using independent SuShiE models from TOPMed MESA, the NHW effect direction validated (*Z*_*NHW*_ = -2.513). No genome-wide significant GWAS SNPs were present within ±1 Mb (*P*< 5 * 10 ^−8^; the most significant association is from rs1072218 *P* = 5.50 * 10 ^− 5^). The direction agrees with NHW AD TWAS in whole blood, conducted by Sun et al.^56^ (*oR* = 0.97; 0.96-0.99).

*COG4* showed a significant association with AD risk in NHW (*q*_*c*_ < 0.05, *Z*_*NHW*_ = 4.601, *P* = 4.20 * 10^−6^ ; population-specific *PIP* > 0.8). SuShiE fine-mapping identified 9 SNPs in two credible sets, with strong anticorrelation of effects between NHW and the other populations 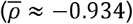. Expression was genetically predictable in both NHW (*R*^2^ = 0.0487, *P* < 0.05) and AA (*R*^2^ = 0.123, *P*< 0.05), yet AA showed no TWAS association (*Z*_*AA*_ = -0.00317, *P*= 0.997). No genome-wide significant GWAS SNPs were present within ±1 Mb (*P*< 5 * 10^−8^; the most significant association is from rs3752786 *P* = 3.98 * 10^−6^). Using independent SuShiE models from TOPMed MESA, the NHW direction replicated but was not significant (*Z*_*NHW*_ = 0.281, *P* = 0.779); attenuation is plausibly due to reduced cis-SNP coverage, differing fine-mapped eQTLs in MESA (±500 kb; MAGENTA *N*_*snp*_ = 1358 vs. MESA *N*_*snp*_ = 783, -42.3%) and potential age/AD-context differences.

In HISP TWAS, one gene reached significance at *q*_*c*_ < 0.05: *GARIN2* (also known as *FAM71D*) on 14q23.3 (*Z*_*HISP*_ = −4.622, *P* = 3.80 * 10^−6^) (Figure 3 (c)). No SuShiE credible sets were identified and 5-fold CV was not significant (*R*^2^ = −0.00345, *P* = 0.991); SuShiE posterior weights in HISP were uniformly small (max magnitude ≈ 1* 10 ^− 5^), indicating the TWAS signal is likely to be confounded by horizontal pleiotropy. The result may instead reflect aggregated weak SNP effects within HISP that may tag other variants in the region (Figure S10).

In AA TWAS, *MARK4* was significant (*Z*_*AA*_ = −4.710, P= 2.50 x 10 ^−6^) within the *APOE* region (∼ 170,000 bp from *APOE*). SuShiE fine-mapping identified 13 SNPs across five credible sets; CS1 showed near-perfect cross-ancestry concordance 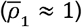, while averaging across all sets indicated heterogeneity 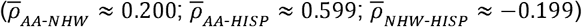. Gene expression predictive performance was not significant in AA (*R*^2^ = 0.0033; P = 0.190), but was significant in NHW and HISP 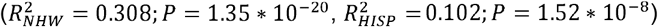. In TOPMed MESA, the AA effect direction was consistent but not significant (*Z*_*AA*_ = −1.662, *P* = 9.66 * 10^−2^). Given regional complexity and proximity to *APOE* (Figure S9), this association should be interpreted with caution.

We explored shared TWAS associations using Inverse-Variance Weighted (IVW) Meta-analysis. There are 23 statistically significant (*q*_*c*_ < 0.05; *P*_*META*_ < 1.4 * 10^−4^) genes, and 8 of them had MA-FOCUS’s PIP > 0.8 (Figure 3 (d)). All fine-mapped meta-analysis associations were driven mostly by the NHW population, due to its larger GWAS sample size. Relative to the NHW TWAS, no new associations became significant after meta-analysis. However, the sign concordance of the 8 identified genes between NHW and the other two populations is 62.5%, which demonstrates moderate consistency and some heterogeneity across populations.

In total, we identified multiple significant TWAS associations across chromosomes 2, 8, 11, and 19, with variable patterns of genetic regulation across populations (Table 2).

**Table 2:**
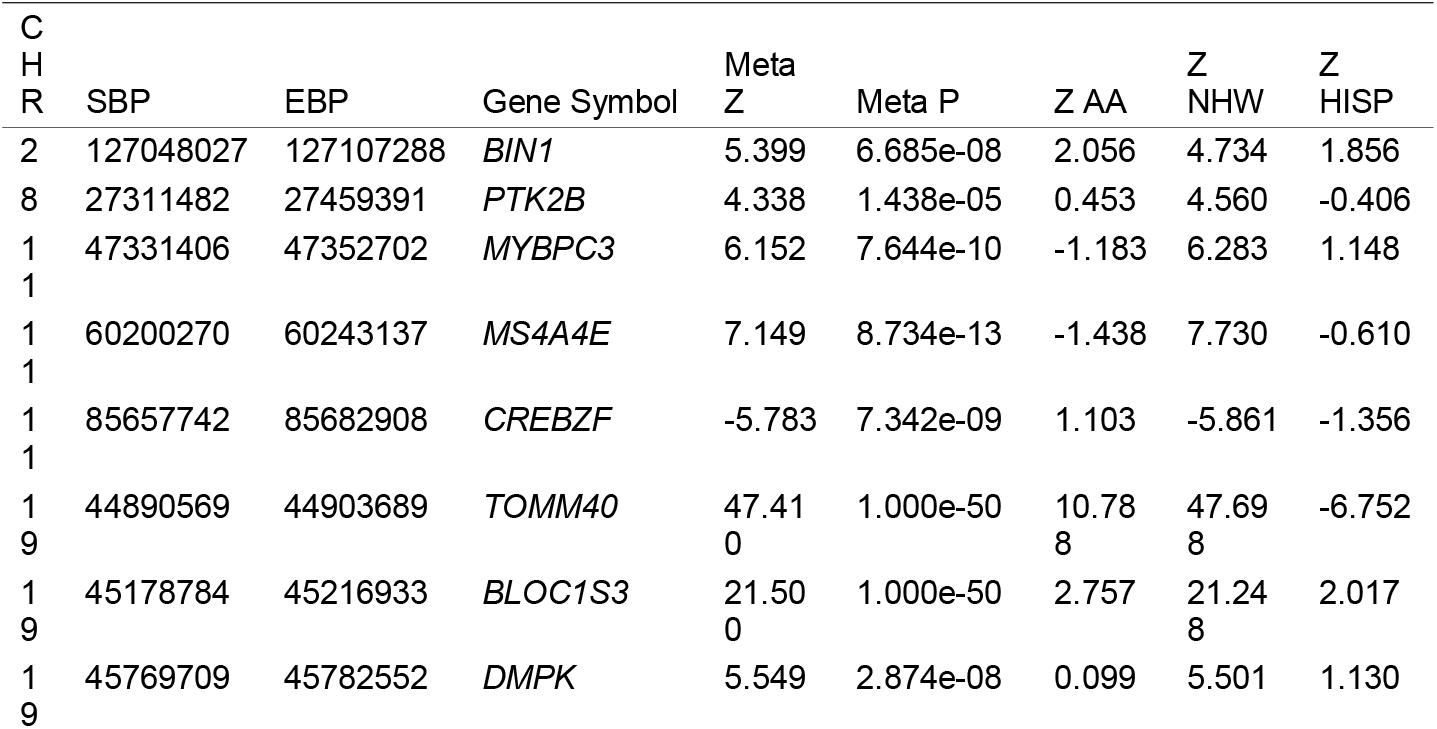
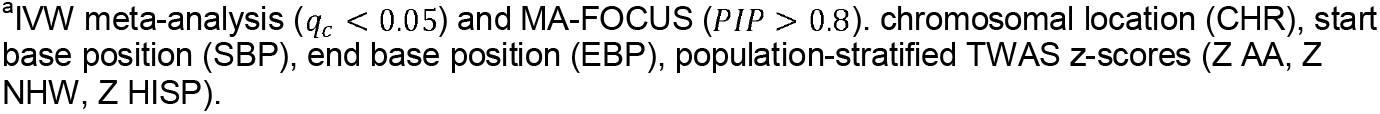
Multi-population TWAS significant genes identified by IVW meta-analysis.

#### Chromosome 2 - *BIN1*

*BIN1* showed highly consistent TWAS associations across AA, NHW, and HISP populations, with fine-mapping implicating three SNPs in two credible sets (mean cross-population effect size correlation 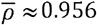). The lead eQTL rs11682128 (previously linked to *BIN1* expression in blood and brain^57^) was in moderate LD (*r*^2^ ≈ 0.34; 53,336 bp away) with the top GWAS SNP rs6733839^4^. 5-fold cross-validation *R*^2^ (0.116-0.263) was significant in all populations.

#### Chromosome 8 - *PTK2B*

*PTK2B* was significant in the meta-analysis and fine-mapping identified 28 SNPs in three credible sets with high effect size concordance 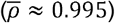. Notably, the 28 SNPs are in minimal LD with the GWAS index SNP^4^, rs2741342, with the highest LD in the first credible set (*r*^2^ ≈ 0.1). Predictive performance (*R*^2^ = 0.040 - 0.120) was significant in all populations, though TWAS significance was observed mainly in NHW, with consistent direction across datasets.

#### Chromosome 11 - *MYBPC3, MS4A4E*, and *CREBZF*

*MYBPC3* showed the strongest signal in NHW, with high cross-population effect size correlation 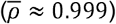. *R*^2^ was significant in only NHW (*h*^2^ = 0.052). *MS4A4E* was driven by the NHW signal, with weak or negative effect size correlation with AA and HISP (AA-NHW: ρ ≈ 0.004; AA-HISP: ρ ≈ −0.004) and significant predictive performance in AA. *CREBZF* showed no significant *R*^2^ (*P* > 0.05), or fine-mapped variants, and was likely influenced by nearby strong GWAS signals (*P*< 1 * 10 ^− 8^ from 86Mb to 86.2Mb).

#### Chromosome 19 - *TOMM40, DMPK*, and *BLOC1S3*

Among the three genes near the *APOE* locus, only the SuShiE TWAS prediction model for *TOMM40* contained the *APOE*-ε4 allele, rs429358^4^ in its first CS.

*TOMM40* exhibited strong but heterogeneous associations, with significant *R*^2^ only in NHW and variable effect directions across populations. The learned prior effect size correlations showed substantial heterogeneity across populations especially in HISP, with moderate positive correlation between AA and NHW 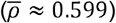, but negative correlations between AA-HISP 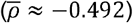 and NHW-HISP 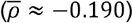. *DMPK* showed consistent positive direction across populations, moderate cross-population correlations 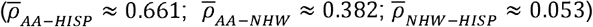, and significant predictive performance (*R*^2^ = 0.026 − 0.146) in all groups. *BLOC1S3* had highly consistent effect sizes across populations 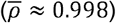 but significant *R*^2^ only in NHW (*R*^2^ = 0.031; *h*^2^ = 0.048).

Across loci, patterns emerged where some genes (*BIN1, PTK2B, BLOC1S3*) had consistent cross-population genetic regulation, whereas others (*MYBPC3, TOMM40, MS4A4E*) displayed marked population-specific architectures in one group. Using direction as the primary criterion, we observed replicated effects in TOPMed MESA for several discovery genes in at least two ancestries-notably *BIN1, PTK2B, DMPK, TOMM40*, and *MYBPC3*, and *MS4A4E*-with significance most evident in NHW and generally insignificant in AA and HISP, consistent with GWAS sample size differences. To assess prediction model quality, we evaluated cross-validation R^2^ for each gene-population pair Table S9. Our prioritized genes - *BIN1, PTK2B, DMPK, COG4* - showed significant R^2^ in the populations where TWAS associations were detected. Two nominally significant associations (*GARIN2, CREBZF*) lacked significant h^2^ and R^2^ in any population and are likely confounded by horizontal pleiotropy. Detailed association results, cis-*h*^2^, *R*^2^, and replication outcomes are presented in Table 2, Table S9, and Table S11.

### Sparse models using fine-mapped eQTLs capture key TWAS associations

Across fine-mapped TWAS hits (MA-FOCUS PIP>0.8), SuShiE eQTL credible sets typically did not include the GWAS index SNPs. At *BIN1*, the top CS SNP rs11682128 was only modestly correlated with the NHW index SNP rs6733839 (*r*^2^ ≈ 0.34). Similar patterns were observed at *PTK2B* and *DMPK*, where rs2741342 and rs429358 are not in the credible sets for each gene respectively. However, in the *APOE* region *TOMM40*’s CS1 included the ε4-defining rs429358 and drove heterogeneous associations across populations. Dense-versus-sparse comparisons showed that restricting models to credible-set variants largely preserved association magnitudes (Table 3; Figure S6), supporting a fine-mapped variant-level regulatory basis for these TWAS signals.

**Table 3:**
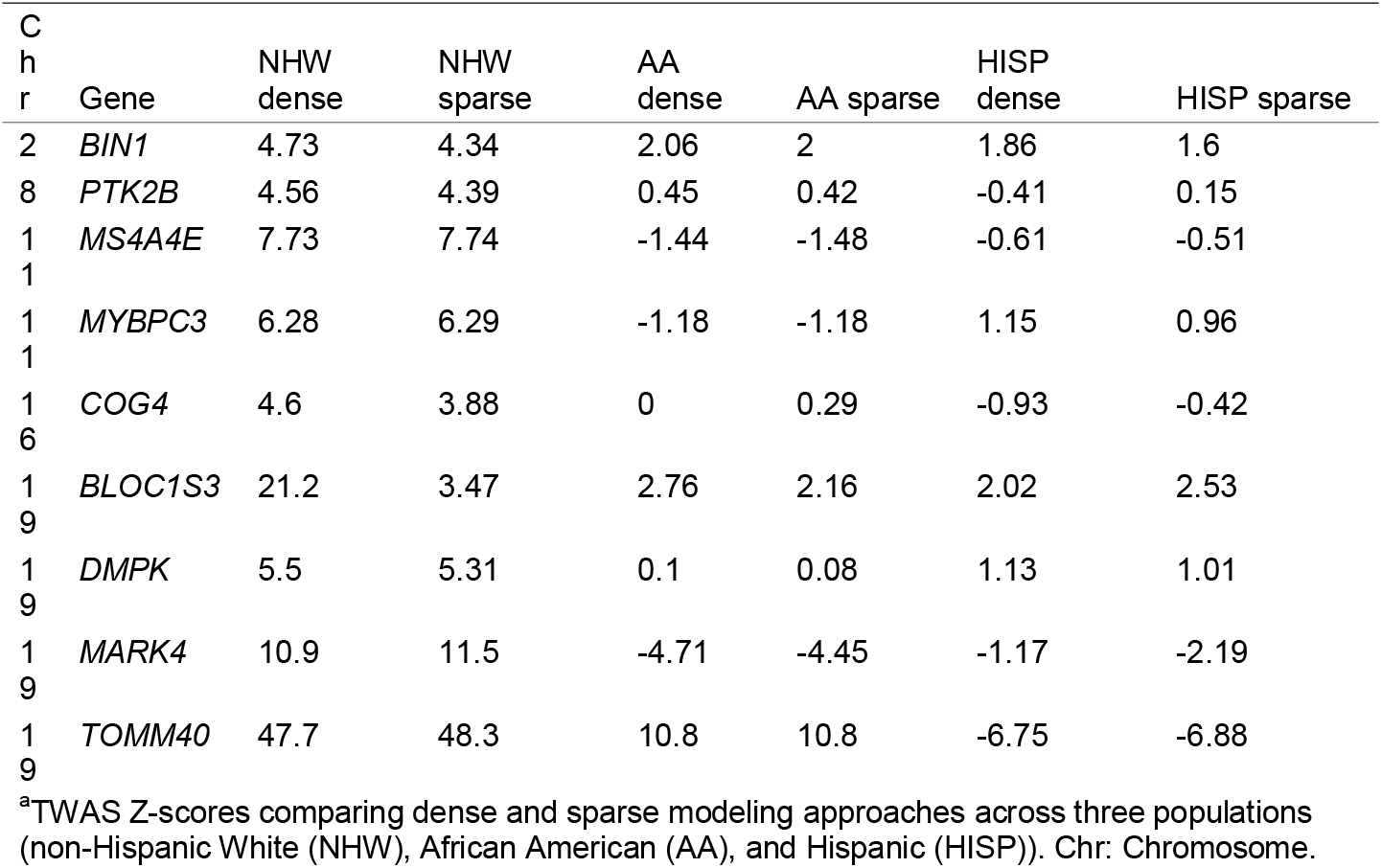
Comparison of Dense and Sparse TWAS Z-scores for Dense Model-Identified Significant Genes.

To evaluate whether restricting prediction models to fine-mapped credible-set variants preserves TWAS signals, we compared TWAS Z-scores obtained from dense SuShiE models (all cis-SNPs within ±500 kb) to those from sparse models restricted to credible-set SNPs. Across genes with at least one credible set, dense and sparse TWAS Z-scores were highly concordant in all three populations (AA: r = 0.851, n = 8,884; NHW: r = 0.869, n = 8,921; HISP: r = 0.828, n = 8,697; Figure S6), indicating that credible-set variants capture most of the association information present in the full cis-region models. Consistent with this, cross-validated predictive R^2^ was comparable between dense and sparse models across all populations (Table S12), demonstrating that fine-mapped variants capture the predictive signal without requiring all cis-SNPs. We further summarized fine-mapping characteristics for genes highlighted in the sparse-model analyses, including credible set sizes and posterior inclusion probabilities, in the Supplementary Material (Table S8).

The comparison revealed remarkable consistency between dense and sparse models for most genes (Table 3). Key AD-associated genes like *BIN1, TOMM40, COG4*, and *MARK4* showed particularly stable associations across both approaches. For instance, *BIN1* maintained consistent negative associations across all populations in dense and sparse models (NHW: 4.73 vs 4.34; AA: 2.06 vs 2.00; HISP: 1.86 vs 1.60), with only slight attenuation in the sparse model. Similarly, *TOMM40* preserved its strong population-specific pattern, showing nearly identical Z-scores in both approaches for all populations (NHW: 47.7 vs 48.3; AA: 10.8 vs 10.8; HISP: -6.75 vs -6.88). However, the strong and heterogeneous associations of *TOMM40* are likely driven by the APOE effect, as the APOE ε4 allele, rs429358, is present in the first CS.

The candidate gene association we identified, *COG4*, also demonstrated stability between dense and sparse models (NHW: 4.6 vs 3.88; AA: 0 vs 0.29; HISP: - 0.93 vs -0.42), suggesting that its association with AD is primarily driven by the fine-mapped eQTLs.

However, some genes showed notable differences between the two approaches. For example, *BLOC1S3* in the APOE region showed different magnitudes of association in the dense versus sparse models in the NHW population (21.2 vs 3.47). This difference could be explained by the complex LD structure and APOE signals in the region, where many variants could contribute to the association signal through LD with known AD risk variants.

Overall, our comparison demonstrates that fine-mapped eQTLs in credible sets can largely explain the TWAS associations for most genes, particularly those with strong and consistent effects across populations. The stability of associations between dense and sparse models provides additional support for the sparse genetic architecture of gene expression regulation and validates our fine-mapping approach.

As illustrated in Figure 4, the variant rs11682128 in *BIN1* (CS1) exhibited consistently positive eQTL weights across all three populations, with magnitudes ranging from 0.44 to 0.64. The visualization further revealed population-specific patterns in the corresponding GWAS signals, with the strongest Z-scores observed in the NHW population (∼4.5), followed by more moderate signals in AA and Hispanic populations (1-2), which may explain the magnitude of TWAS associations across ancestries. The difference could be due to the sample size differences in GWAS studies, or real effect size differences across populations as we observe with APOE^5^. Despite these differences in GWAS signal intensity, the direction and relative magnitude of the TWAS associations remain consistent (Table 3), suggesting that the identified credible set variants captured the core regulatory mechanisms influencing *BIN1* expression in relation to AD risk. The genomic context provided in the bottom panel demonstrates that the two credible sets are located in the intronic region of *BIN1*. Collectively, this integrated view of *BIN1* regulation supports our finding that sparse models using only credible set variants can effectively recapitulate the signals detected in dense models, while providing greater interpretability regarding the specific variants driving these associations.

**Figure 4:**
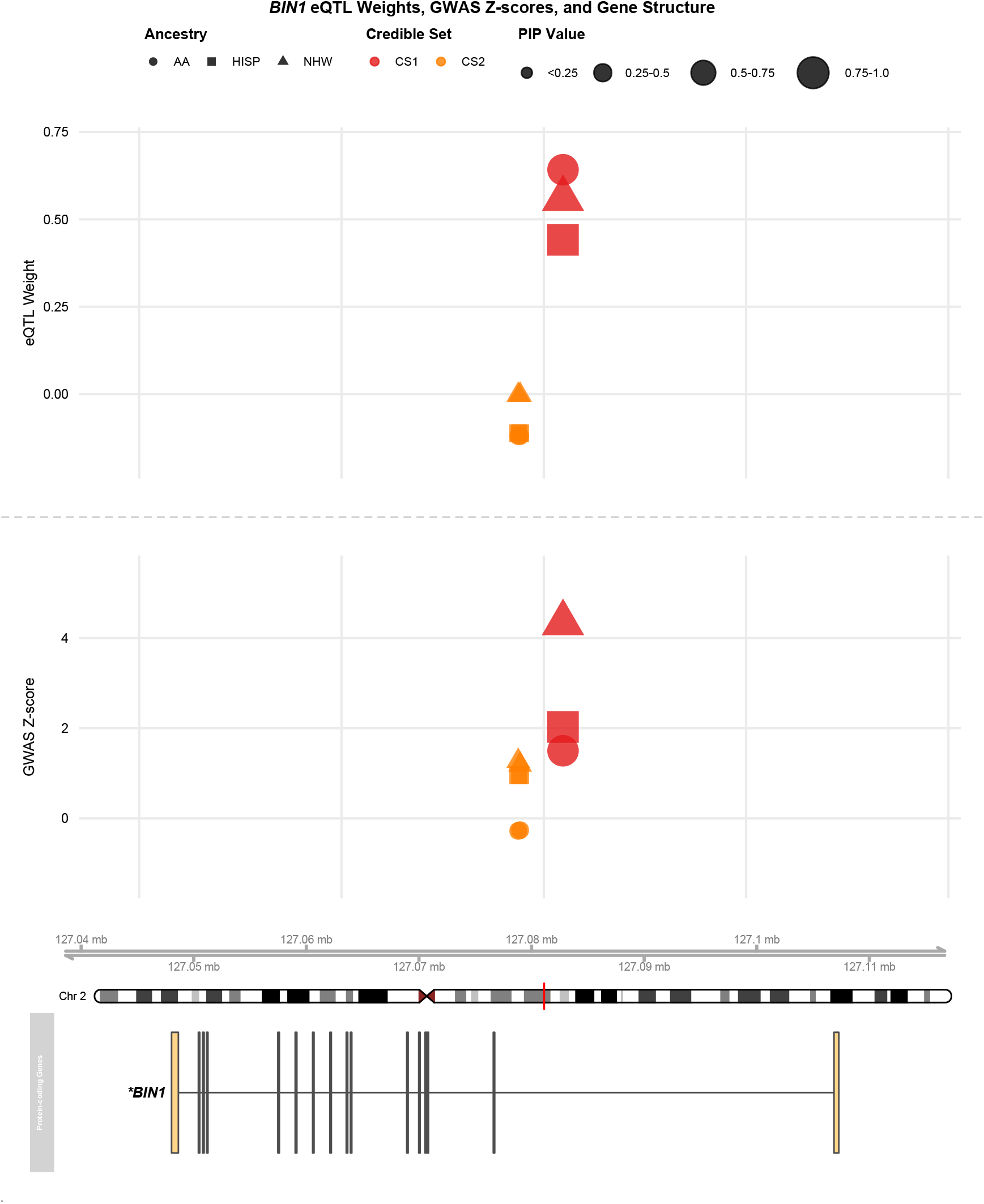
Fine-mapped eQTLs and GWAS associations for the *BIN1* locus. The figure displays the relationship between eQTL weights (top panel), GWAS Z-scores (middle panel), and gene structure (bottom panel) for *BIN1* on chromosome 2. In the top and middle panel, eQTL weights and GWAS associations are shown across three ancestral populations (African American (AA, circles), Hispanic (HISP, squares), and non-Hispanic White (NHW, triangles)), with variants colored by credible set membership. Point sizes indicate posterior inclusion probability (PIP) values. The bottom panel illustrates the genomic structure of *BIN1*. In CS1, there is only one variant, rs11682128; In CS2, there are two variants, rs6714626 and rs6710752.

*COG4* is identified as an AD-related gene from our main analysis. When we focused on the sparse model, which included only 9 variants in 2 credible sets, we observed a similar TWAS association (*Z*_*NHW*_ = 3.88) in the NHW population, while the other two populations did not show significant associations (Table 3). After analyzing the SNP weights and GWAS associations for these fine-mapped SNPs (Figure 5), we noticed substantial differences in the NHW population, particularly in the second credible set. For example, rs11639579, the fine-mapped SNP in the second credible set with the largest PIP (*PIP* = 0.709), exhibited the largest difference in NHW’s GWAS association and SNP weight compared to the other two populations (Table 3). We observed that this variant is also located near *IL34*, but the SNP is not fine-mapped in *IL34*’s credible set, and the TWAS association of *IL34* is not significant in the dense model.

**Figure 5:**
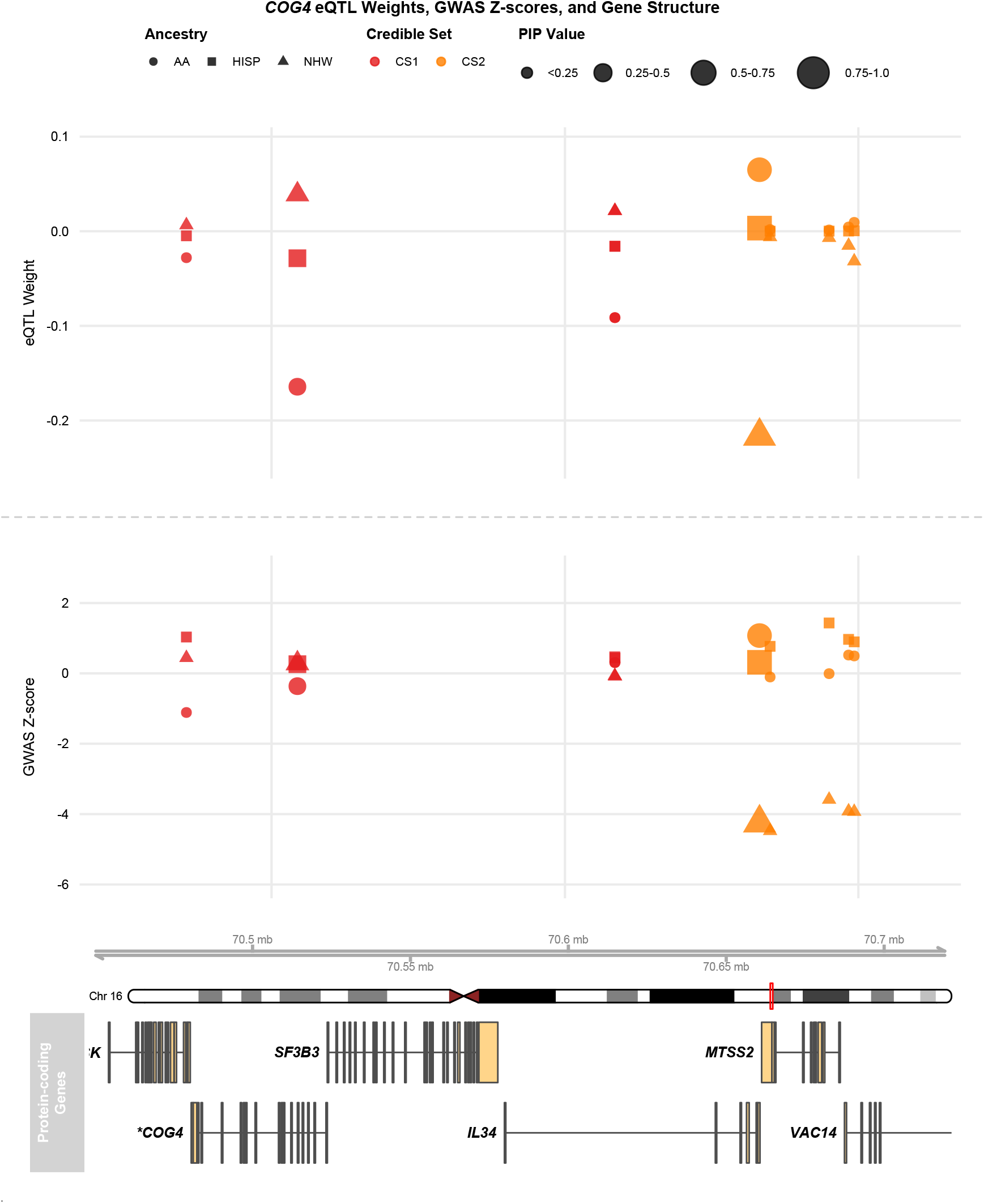
Fine-mapped eQTLs and GWAS associations for the *COG4* locus. The figure displays the relationship between eQTL weights (top panel), GWAS Z-scores (middle panel), and gene structure (bottom panel) for *COG4* on chromosome 16. In the top and middle panels, eQTL weights and GWAS associations are shown across three ancestral populations (African American (AA, circles), Hispanic (HISP, squares), and non-Hispanic White (NHW, triangles)), with variants colored by credible set membership. Point sizes indicate posterior inclusion probability (PIP) values. The bottom panel illustrates the genomic structure of *COG4*.

To characterize the regulatory mechanism underlying the the variants in the second credible set of *COG4*, we queried enhancer chromatin states and chromatin interaction data from FILER (Tables S14-S16). The lead variant chr16:70666311_A_C overlapped an active enhancer chromatin state (EnhA2) in GM19238 lymphoblastoid cells based on EpiMap ChromHMM annotations. This enhancer annotation supports a distal regulatory mechanism whereby the variant affects *COG4* expression by modulating enhancer activity.

We also examined promoter-capture Hi-C (pcHi-C) data from primary hematopoietic cells as supplementary evidence for physical chromatin contact. The lead variant is located within a genomic region (chr16:70,652,543-70,673,594) that shows detectable, though sub-threshold, chromatin contact with the *COG4* promoter (chr16:70,522,638-70,524,465) (max CHiCAGO score = 2.65 in macrophages; significance threshold ≥5).

The remaining CS2 variants showed extensive enhancer chromatin states across blood cell types. chr16:70696670_A_C exhibited the broadest enhancer annotation, overlapping active enhancer states (EnhA1) in 21 blood cell types.

Together, the strong eQTL fine-mapping signal (*PIP* = 0.71) combined with active enhancer chromatin states supports chr16:70666311_A_C as the likely causal variant regulating *COG4* expression through a distal enhancer mechanism.

## Discussion

In this multi-ancestry TWAS of AD, we show that jointly modeling cis-eQTLs across African American, non-Hispanic White, and Hispanic groups improves fine-mapping resolution and yields more interpretable, variant-resolved associations. Through the lens of whole-blood gene expression, we implicate regulatory variants beyond GWAS index SNPs at established loci at *BIN1, PTK2B*, and *DMPK*. We further nominate *COG4* as a candidate gene for AD.

Importantly, we demonstrate that sparse credible-set models from our fine-mapping recapitulate dense-model associations, concentrating evidence onto compact variant sets without a significant loss of association signal. Our multi-ancestry fine-mapping of eQTL results decreased the number of SNPs in each credible set while also increasing the overall number of credible sets. These findings are consistent with the results reported in Lu et al.^31^ and provide additional empirical evidence supporting the sparse genetic architecture of gene expression regulation previously reported in multiple studies^20–22^. Our results also align with prior studies showing that while differences in LD and allele frequencies across ancestral groups might lead to heterogeneous eQTL effects, the correlation is largely consistent across populations^31,58–60^. However, we observed lower cross-ancestry effect correlations compared to Lu et al.^31^, with approximately 30% of genes showing anti-correlated effects-predominantly reflecting opposing direction effects. This discrepancy likely stems from the heterogeneous cellular composition of whole blood and a lower fraction of significantly heritable eGenes in MAGENTA (49.7% vs. 89.4% in at least one population), which introduces greater noise into effect estimates. Restricting to genes with significant cis-heritability substantially improved concordance, consistent with the pattern reported by Lu et al^31^.

Our multi-population TWAS analysis revealed both established and additional candidate gene associations with AD risk. Because TWAS tests associations between disease risk and genetically predicted expression, the signal is expected to be driven by cis-regulatory instruments rather than necessarily the GWAS index SNP. The contribution of our multi-ancestry SuShiE analysis is that it fine-maps these instruments into credible sets, enabling direct comparison between high-PIP eQTL variants and published GWAS sentinel variants. Across prioritized loci (*BIN1, PTK2B*, and *DMPK*), the credible-set variants were often only weakly correlated with the GWAS index SNPs, highlighting additional candidate functional variants within established AD loci. *BIN1* showed consistent TWAS effects and was directionally supported using TOPMed MESA models across ancestries; rs11682128 exhibited shared eQTL effects across all three populations as previously reported^57^ in NHW population alone. We also identified two additional variants in a second credible set, suggesting the existence of more than one cis-regulatory signal contributing to *BIN1* expression.

We also identified *CEBPZOS* (that has been reported from prior TWAS^57^) and *COG4*, neither of which have nearby genome-wide significant GWAS signals. To the best of our knowledge, *COG4* had not been identified as an AD-related gene previously. Notably, single-ancestry fine-mapping in NHW alone identified no credible sets, whereas including all three populations yielded two credible sets, underscoring the advantage of multi-ancestry fine-mapping for variable selection. Specifically, four variants in the first credible set had higher allele frequency in AA than the other two populations (Table S13). *COG4* encodes a component of an oligomeric protein complex that plays a crucial role in the structure and function of the Golgi apparatus, which is essential for processing and distributing proteins necessary for neuronal function. Notably, rs11639579, the SNP in the second credible set with the largest PIP (*PIP* = 0.709), exhibited substantial differences in GWAS association and SNP weight in the NHW (-0.217; negative) population compared to AA (0.065; positive) and HISP (0.003; near-zero) populations. Additionally, rs11639579 resides within an active enhancer element approximately 140 kb from the *COG4* promoter; while pcHi-C data showed sub-threshold chromatin contact (CHiCAGO = 2.65), the strong eQTL association (PIP = 0.71) provides evidences for distal regulation of *COG4* expression.

According to Agora^61^, an AD knowledge portal, *COG4* has meaningful expression across brain regions. The differential expression, between AD cases and controls, is significant in cerebellum (log_2_*FC* = -0.149, *P*= 1.44 * 10 ^− 3^) and parahippocampal gyrus (log_2_*FC* = 0.0827, *P* = 1.44 * 10 ^− 2^). Furthermore, Golgi fragmentation occurs in many neurodegenerative diseases. In AD, Golgi fragmentation results in enhanced *APP* trafficking and Aβ production^62,63^. The significant association of genetically predicted *COG4* expression in blood with AD suggests a potentially underexplored link between systemic vesicle trafficking pathways and disease risk.

Previous studies^50–52^ have demonstrated that cis-eQTL effects are largely consistent between disease and control cohorts and healthy individual cohorts, particularly in neuropsychiatric and neurodegenerative disorders. This same consistency was observed in the MAGENTA study for eQTL mapping and TWAS analysis. We chose not to adjust for AD status in our primary eQTL models because, as demonstrated below, eQTL effect sizes are highly consistent regardless of AD adjustment, and statistical adjustment in a case-control design can introduce collider bias. Our sensitivity analyses comparing AD-adjusted and unadjusted models confirmed this approach, showing high concordance in TWAS associations across all populations (*r* ≈ 0.9) (Figure S3). To confirm that concordance holds for the most strongly associated genes, we additionally examined the top 5% of genes by maximum absolute TWAS Z-score.

Concordance remained high in this subset (AA: r=0.943, NHW: r=0.894, HISP: r=0.954). Consistent with this, eQTL effect size correlations between adjusted and unadjusted models were similarly robust in the top 5% subset (AA: r=0.784, NHW: r=0.817, HISP: r=0.863) (Table S6). To further visualize these patterns, we included an additional supplemental figure (Figure S4) in which genes are colored by significance categories, confirming that the global adjusted vs. unadjusted relationship is preserved across ancestries. However, we noted that genes in *APOE* region showed some inflation, especially in NHW population. Among genes with |ΔZ| > 4, 9 of 12 (75.0%) were NHW, 1 (8.3%) were AA, and 2 (16.7%) were HISP; most NHW genes with large ΔZ mapped to the APOE region, while the non-NHW instances did not (Table S7). This confirms that large adjusted-vs-unadjusted divergences are predominantly in NHW’s APOE-region. We observed over 73% overlapping SNPs in credible sets, with the first credible sets, which capture primary regulatory effects, showing particularly high overlap at 75% shared eQTLs (Table S5). The effect size correlations for overlapped eQTLs were remarkably high (*r* ≈ 0.98) (Figure S5). We also compared the TWAS Z-score between AD-adjusted and unadjusted models. Overall, we found most of the significant genes identified in the main analysis were consistent (Table S4). *BLOC1S3* and *TOMM40* in *APOE* region showed 14.2 and 10.9 Z-score reduction after adjusting AD status, but their associations are still significant. Similarly, *MS4A4E* had 2.4 reduction in Z-score after the adjustment. In addition, *CEBPZOS* had 1.55 reduction in Z-score, and did not pass the FDR threshold after adjusting AD status. These findings demonstrate that while preserving disease-related expression patterns, the core genetic regulatory mechanisms remain robust and reliable for the TWAS analysis.

Our study faces several important limitations. First, while whole blood is readily accessible and often used in large-scale genomic studies, it may not optimally reflect the tissue-specific regulatory mechanisms occurring in the brain. Currently, the scarcity of brain-specific eQTL data from diverse populations constrains our ability to conduct optimal TWAS analysis and validate these findings in the most relevant tissue. However, several prior studies observed largely concordant AD TWAS effect size directions and eQTL effect sizes between brain and peripheral tissues^64,65^. Additionally, Huseby et al. found eight fundamental cell biological functions that are altered in whole blood samples from six neurodegenerative diseases^66^, suggesting AD risk can be modulated by multiple systemic factors^67^. Second, the substantial disparity in GWAS sample sizes between populations significantly impacted our ability to detect population-specific associations. This imbalance is reflected in our meta-analysis results, where the significant associations were predominantly driven by NHW signals, as evidenced by the stronger Z-scores in NHW population across most identified genes (Table 2). Nevertheless, despite these power limitations, we observed consistent effect directions across populations for several key genes. Third, a practical limitation of multi-ancestry fine-mapping (including SuShiE) is the requirement to analyze only SNPs shared across populations to ensure matched LD and comparability; such filtering can exclude ancestry-unique variants, biasing credible sets toward shared signals and potentially underestimating truly population-specific regulatory variants, especially in AA and admixed HISP where allele frequencies and LD differ. We attempted to mitigate extreme sparsity by applying a per-population MAC threshold (MAC ≥ 10). Future work should explicitly accommodate ancestry-unique variants in multi-ancestry fine-mapping method developments, and expand recruitment of underrepresented populations to enable adequately powered ancestry-specific fine-mapping and TWAS that are comparable to EUR-based studies. Forth, we acknowledge that reliance on the European-derived LM22 panel may reduce cell type deconvolution accuracy and introduce population-specific bias in our AA and HISP datasets, so these cell-type covariates should be interpreted cautiously and revisited with ancestry-matched or single-cell-derived references as they become available.

Several key future directions could build upon our findings. First, as brain-specific multi-population eQTL datasets become available, replicating our analyses in neural tissue would be valuable for validating these associations and potentially uncovering additional AD-related genes. More critically, ongoing efforts to expand AD GWAS in non-European populations will be crucial for achieving more balanced statistical power across populations. This enhanced power would not only improve our ability to detect population-specific effects but also provide more robust evidence for shared genetic mechanisms across populations. Such advancements would help address the current European bias in genetic studies and contribute to a more comprehensive understanding of AD’s genetic architecture across diverse populations.

## Data and code availability

The analysis code is available at Github (https://github.com/Xinyu-Sun/AD_MA_TWAS_SuShiE_analysis_code). Summary statistics, SuShiE eQTL fine-mapping results, and prediction models generated in this study have been deposited in Zenodo (https://zenodo.org/records/17643736).

## Acknowledgments

This work was supported by grants from the National Institute on Aging R01AG070935 (Bush, Griswold), U19AG074865 (Pericak-Vance, Byrd, Bush, Haines, Kunkle, Reitz, Vance), and RF1AG061351 (Below, Bush, Naj). This work made use of the High Performance Computing Resource in the Core Facility for Advanced Research Computing at Case Western Reserve University.

