## Supplemental Material for "Multi-ancestry Transcriptome-Wide Association Study Reveals Shared and Population-Specific Genetic Effects in Alzheimer’s Disease"

### Supplementary Material

#### Supplemental Figures and Legends

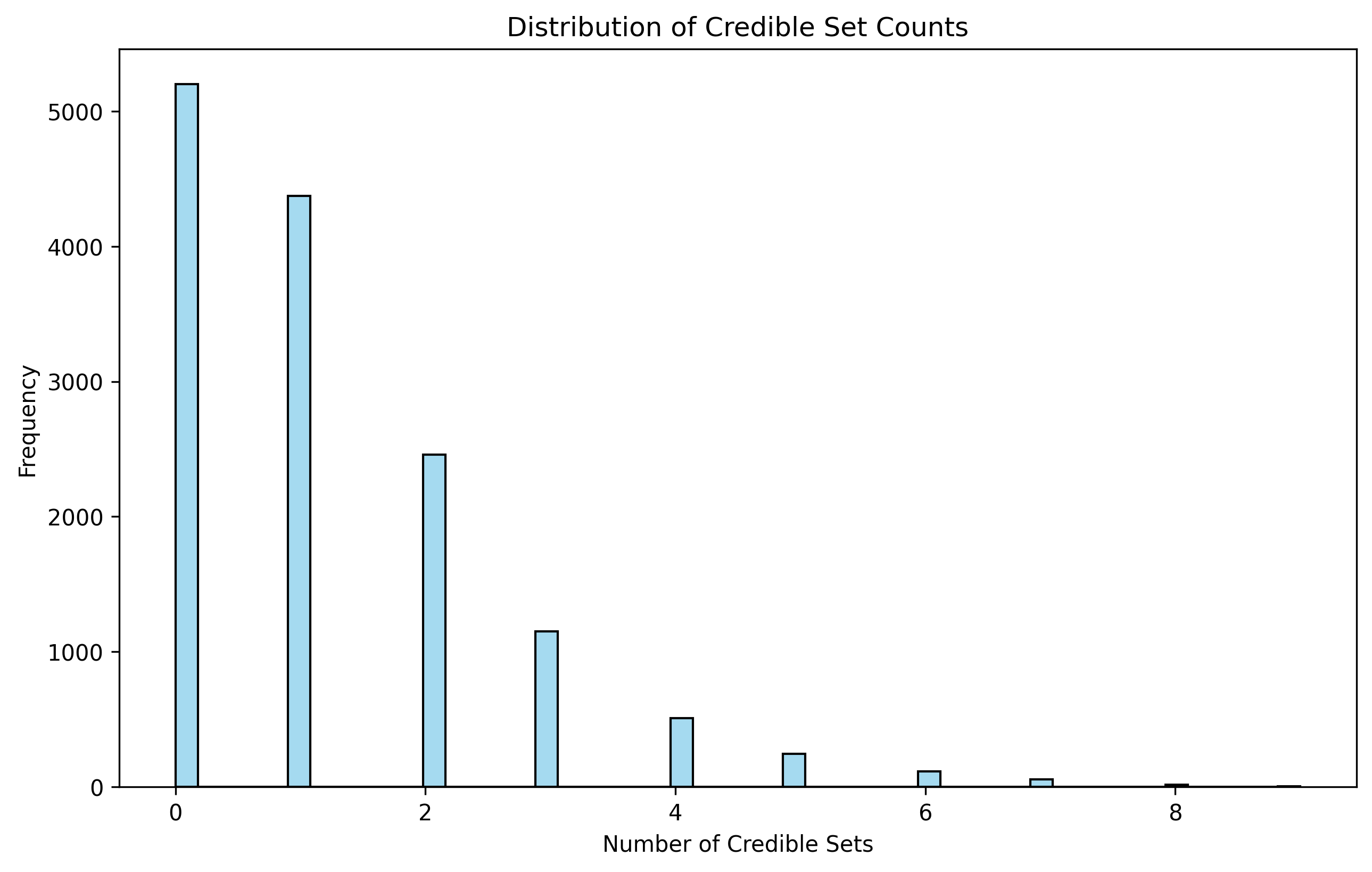

Figure S1: **Distribution of the number of SuShiE credible sets per gene across 14,436 protein-coding genes.** A histogram summarizing the number of credible sets (CS) identified in 14,436 tested protein-coding genes. There are 8,748 genes with at least one CS. The mean number of CS is 1.27, and the median is 1.

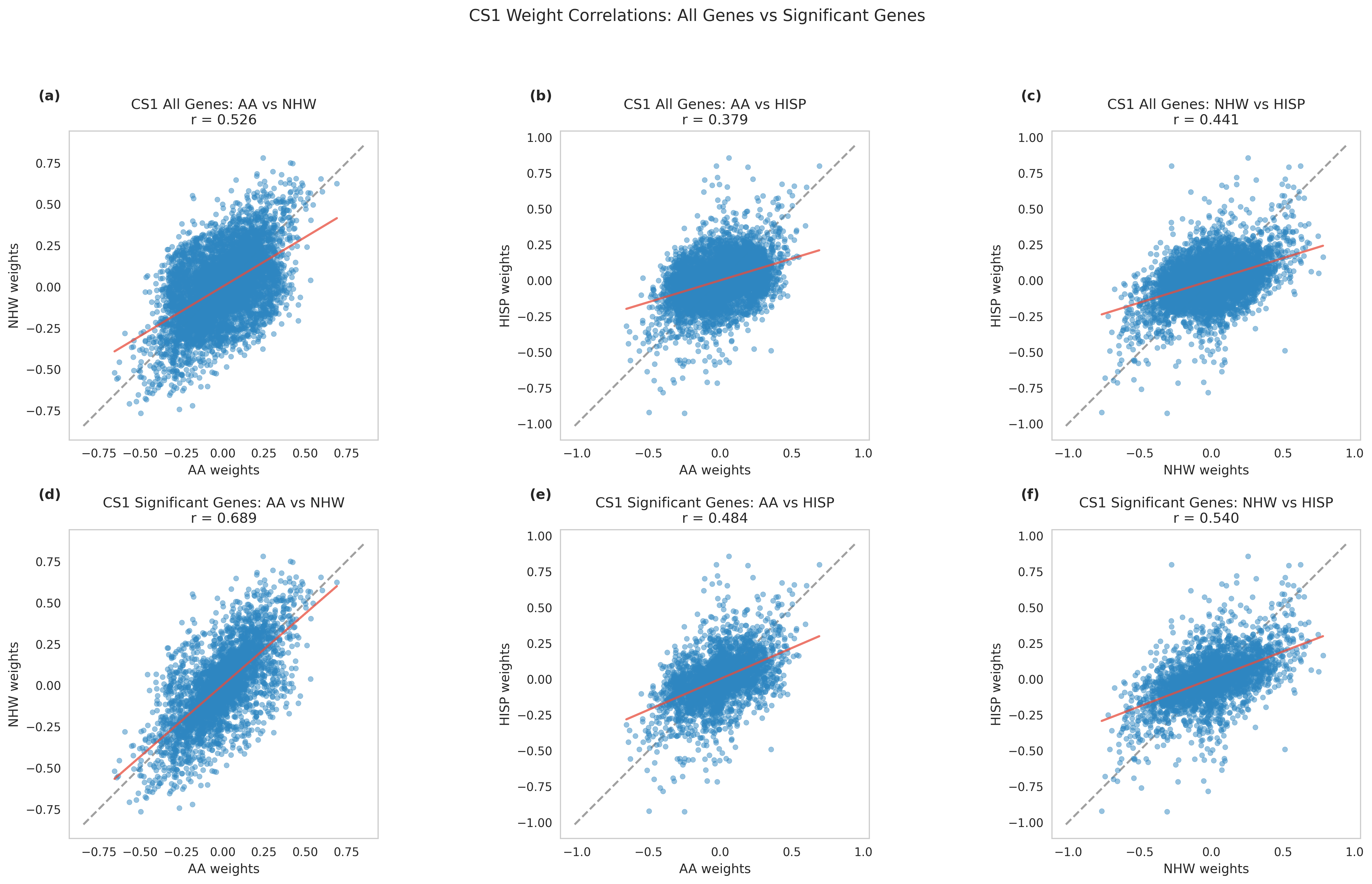

Figure S2: **Pairwise Pearson correlations of SuShiE’s population-specific posterior weights from the first credible set (CS1) across three ancestral populations (AA, NHW, and HISP).** Top row (a-c) shows correlations using all genes, while bottom row (d-f) shows correlations using only genes with significant cis-SNP heritability (P < 0.05) in at least one population. (a,d) Comparison between AA and NHW populations. (b,e) Comparison between AA and HISP populations. (c,f) Comparison between NHW and HISP populations. Gray dashed lines represent the diagonal (y=x), and red solid lines show the best-fit linear regression. Pearson correlation coefficients (r) are shown for each comparison. Higher correlations are observed in significant genes compared to all genes, with the strongest correlation between AA and NHW populations (r=0.689 for significant genes)

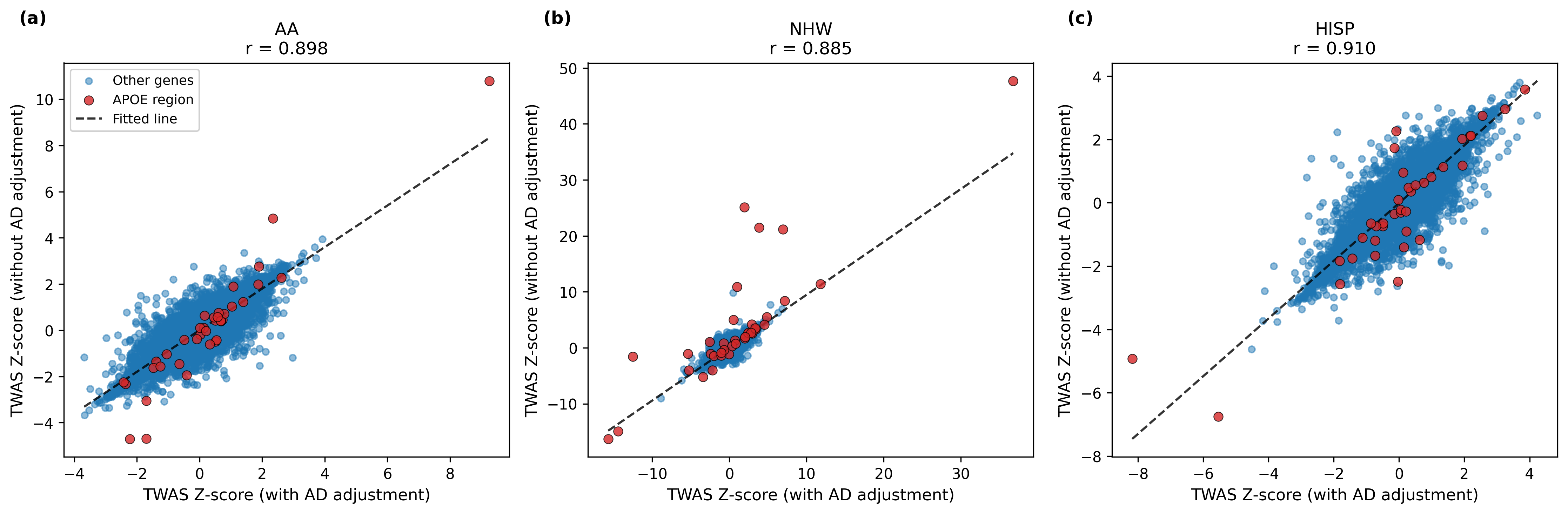

Figure S3: **Correlation of TWAS Z-scores between models with and without Alzheimer’s disease (AD) adjustment across three ancestral populations.** Scatter plots showing the relationship between TWAS Z-scores from models with AD adjustment (x-axis) and without AD adjustment (y-axis) for (a) African American (AA), (b) non-Hispanic White (NHW), and (c) Hispanic (HISP) populations. Genes in the APOE region (chr19: 44.8-45.8 Mb) are highlighted in red; all other genes are shown in blue. The dashed line represents the fitted regression line. Pearson correlation coefficients (r) demonstrate strong concordance between the two models across all populations (AA: r = 0.898, NHW: r = 0.885, HISP: r = 0.910), suggesting that AD adjustment has minimal impact on overall TWAS association patterns while preserving population-specific expression patterns.

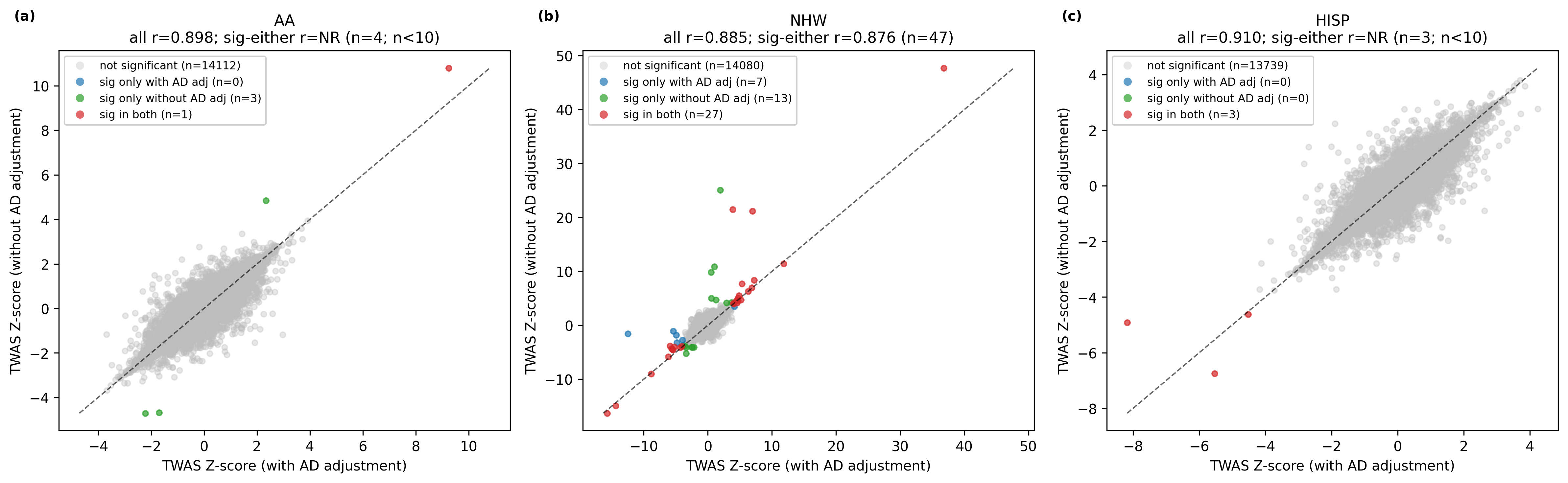

Figure S4: **TWAS Z-score concordance between AD-adjusted and non-adjusted models, colored by significance class across ancestries.** Scatter plots compare TWAS Z-scores from AD-adjusted models (x-axis) versus non-adjusted models (y-axis) for AA, NHW, and HISP analyses. Each point is a gene present in both analyses. Colors indicate FDR-based significance class: grey, not significant in either model; blue, significant only with AD adjustment; green, significant only without AD adjustment; red, significant in both models (FDR < 0.05 within ancestry and model). The dashed black line is the identity line (y = x). Panel titles report Pearson correlation for all genes in each ancestry. Correlation for the significant-gene subset (sig-either) is shown only when the subset size is sufficiently stable (n >= 10). Therefore, sig-either correlations are not reported for AA (n = 4) or HISP (n = 3), but are reported for NHW (n = 47).

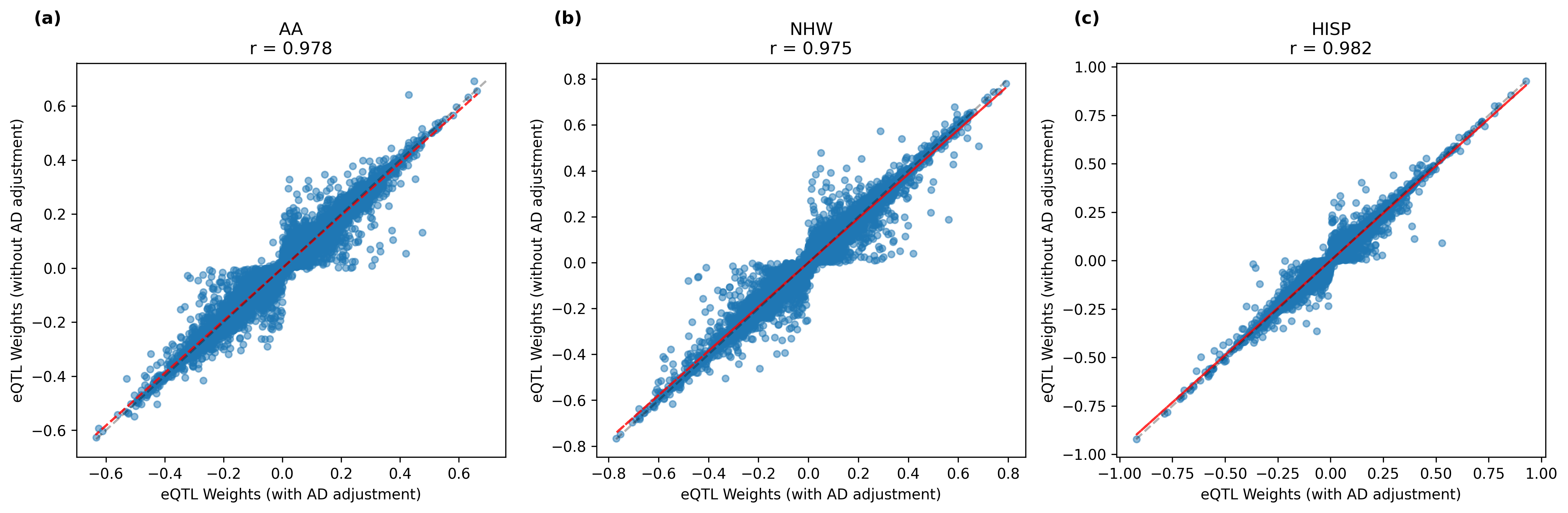

Figure S5: **Comparison of eQTL weights between models with and without Alzheimer’s disease (AD) adjustment across three ancestral populations.** Scatter plots displaying the correlation between eQTL weights from models with AD adjustment (x-axis) and without AD adjustment (y-axis) for (a) African American (AA), (b) non-Hispanic White (NHW), and (c) Hispanic (HISP) populations. Each point represents an individual SNP-gene pair in credible sets. The red dashed lines indicate the best-fit linear regression. Extremely high Pearson correlation coefficients (AA: r = 0.978, NHW: r = 0.975, HISP: r = 0.982) demonstrate that genetic effect estimates remain highly consistent regardless of AD adjustment, suggesting that the underlying genetic architecture of gene expression is robustly captured by both modeling approaches across diverse populations.

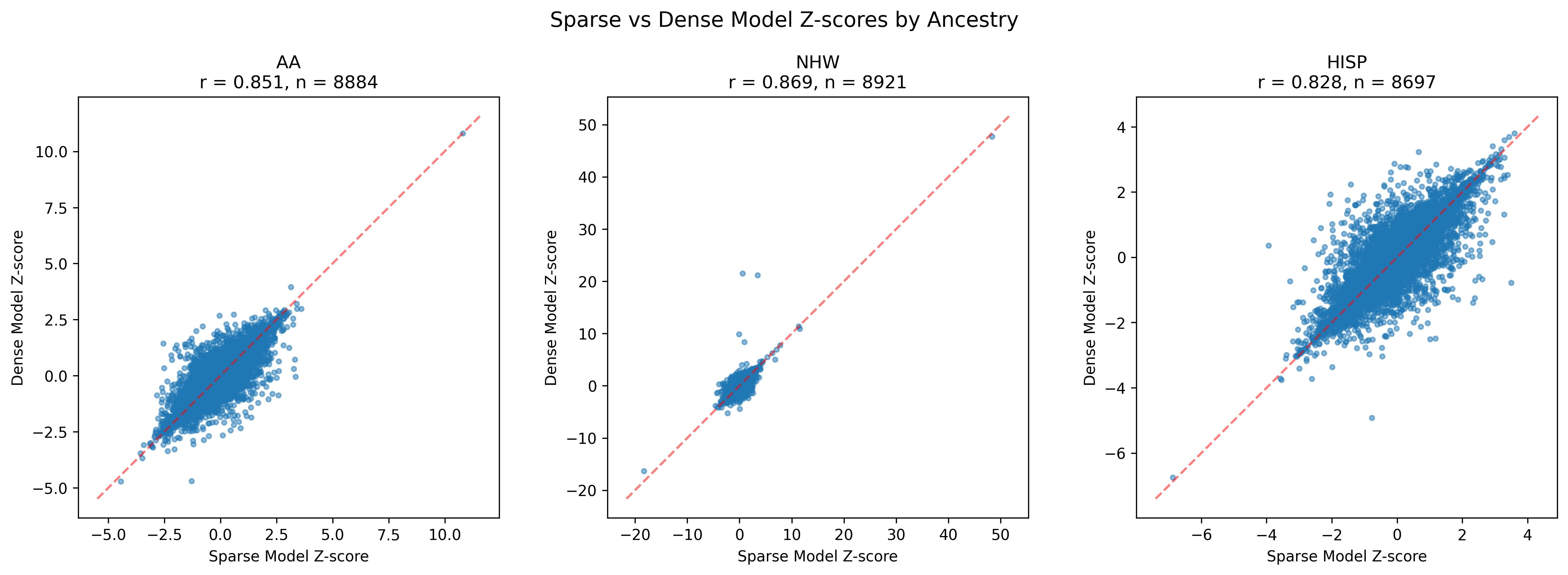

Figure S6: **Correlation of TWAS Z-scores between sparse and dense prediction models across three ancestral populations.** Scatter plots showing the relationship between TWAS Z-scores from sparse models (x-axis) and dense models (y-axis) for (a) African American (AA), (b) non-Hispanic White (NHW), and (c) Hispanic (HISP) populations. Each point represents a gene, and the red dashed line indicates perfect concordance (y = x). Pearson correlation coefficients (r) demonstrate strong agreement between the two model types across all populations (AA: r = 0.851, n = 8884; NHW: r = 0.869, n = 8921; HISP: r = 0.828, n = 8697), indicating that sparse and dense eQTL prediction models produce highly consistent TWAS association estimates regardless of ancestry.

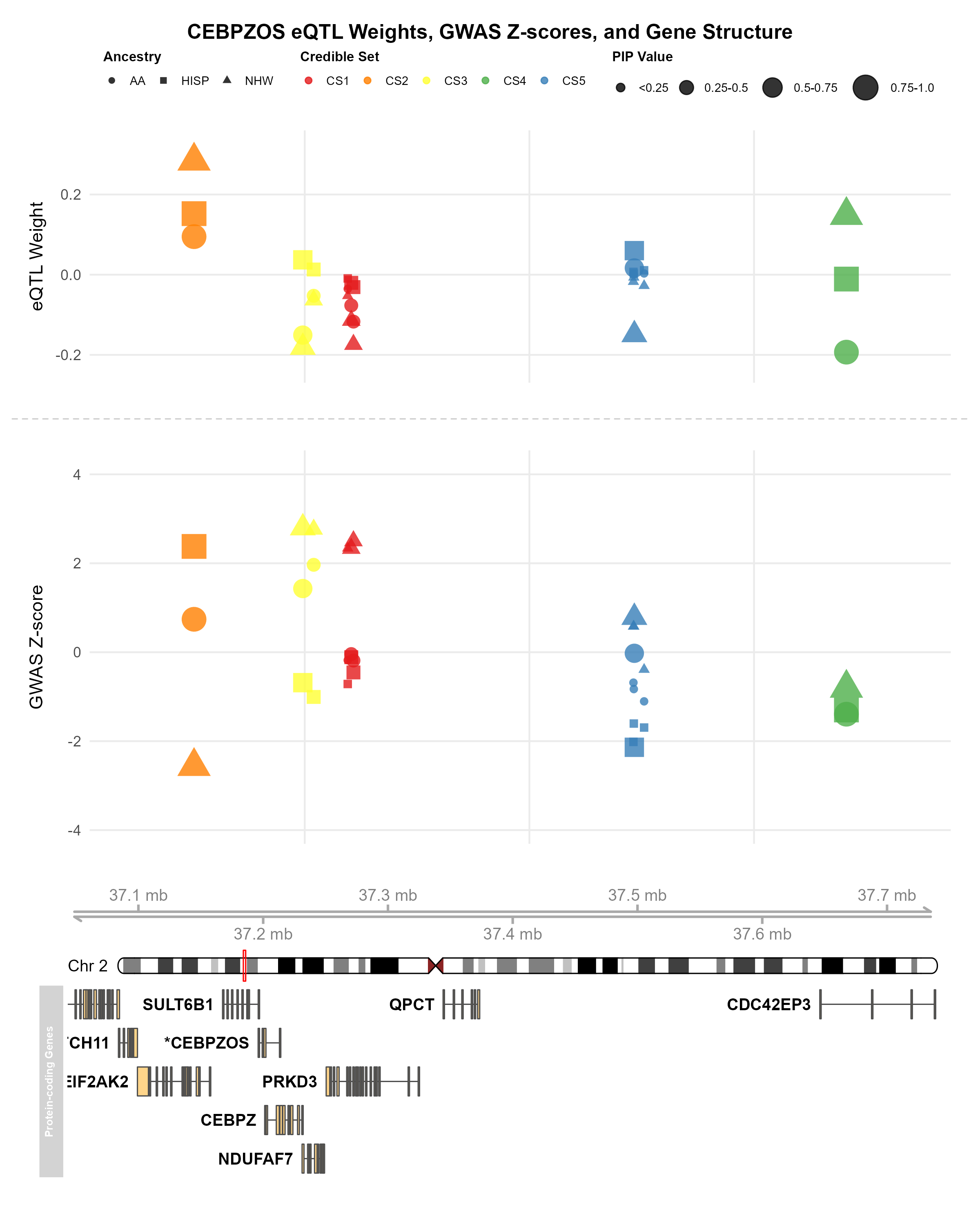

Figure S7: **Fine-mapped eQTLs and GWAS associations for the CEBPZOS locus.** The figure displays the relationship between eQTL weights (top panel), GWAS Z-scores (middle panel), and gene structure (bottom panel) for CEBPZOS on chromosome 2. In the top and middle panels, eQTL weights and GWAS associations are shown across three ancestral populations (African American (AA, circles), Hispanic (HISP, squares), and non-Hispanic White (NHW, triangles)), with variants colored by credible set membership. Point sizes indicate posterior inclusion probability (PIP) values. The bottom panel illustrates the genomic structure of CEBPZOS.

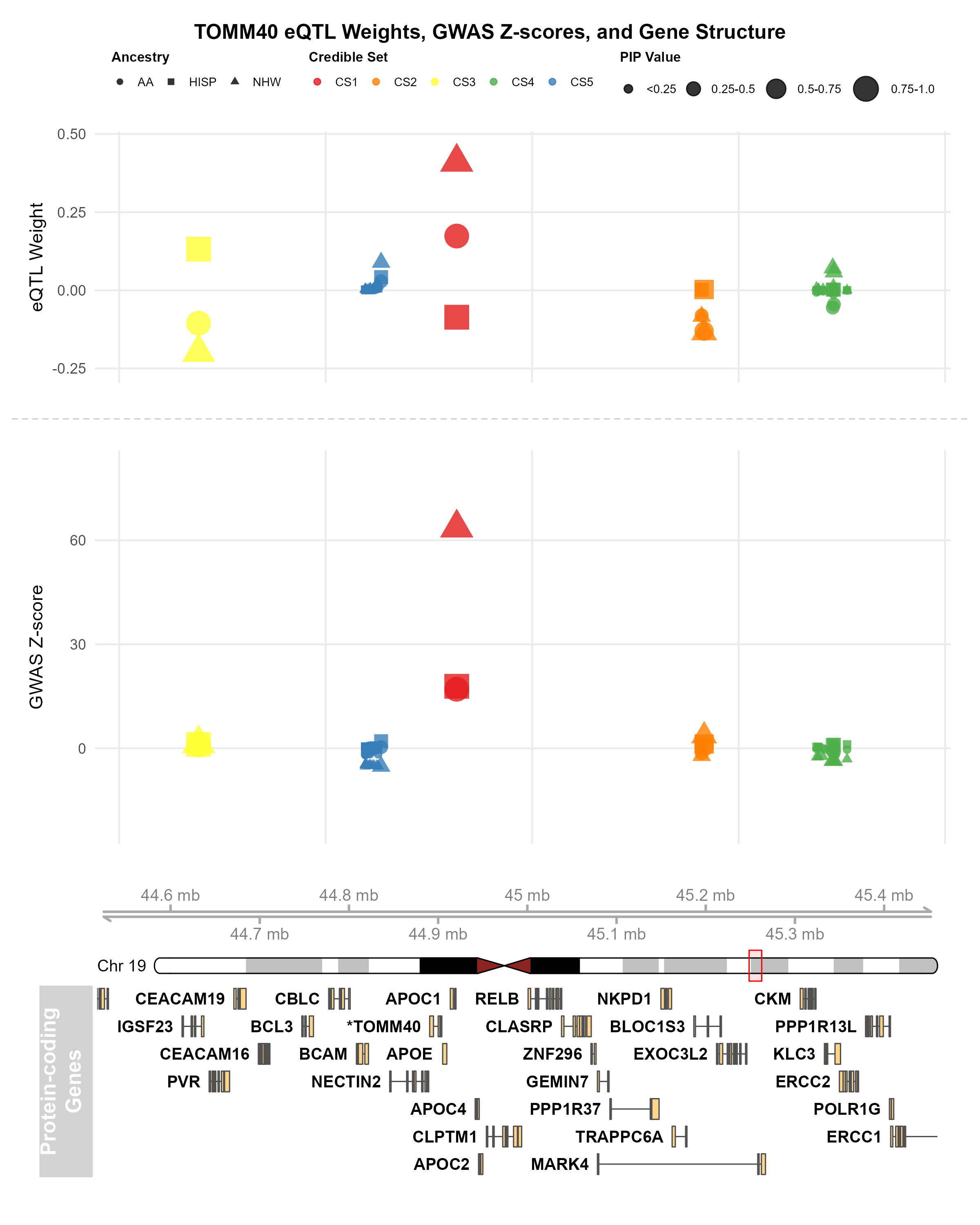

Figure S8: **Fine-mapped eQTLs and GWAS associations for the TOMM40 locus.** The figure displays the relationship between eQTL weights (top panel), GWAS Z-scores (middle panel), and gene structure (bottom panel) for TOMM40 on chromosome 19. In the top and middle panels, eQTL weights and GWAS associations are shown across three ancestral populations (African American (AA, circles), Hispanic (HISP, squares), and non-Hispanic White (NHW, triangles)), with variants colored by credible set membership. Point sizes indicate posterior inclusion probability (PIP) values. The bottom panel illustrates the genomic structure of TOMM40.

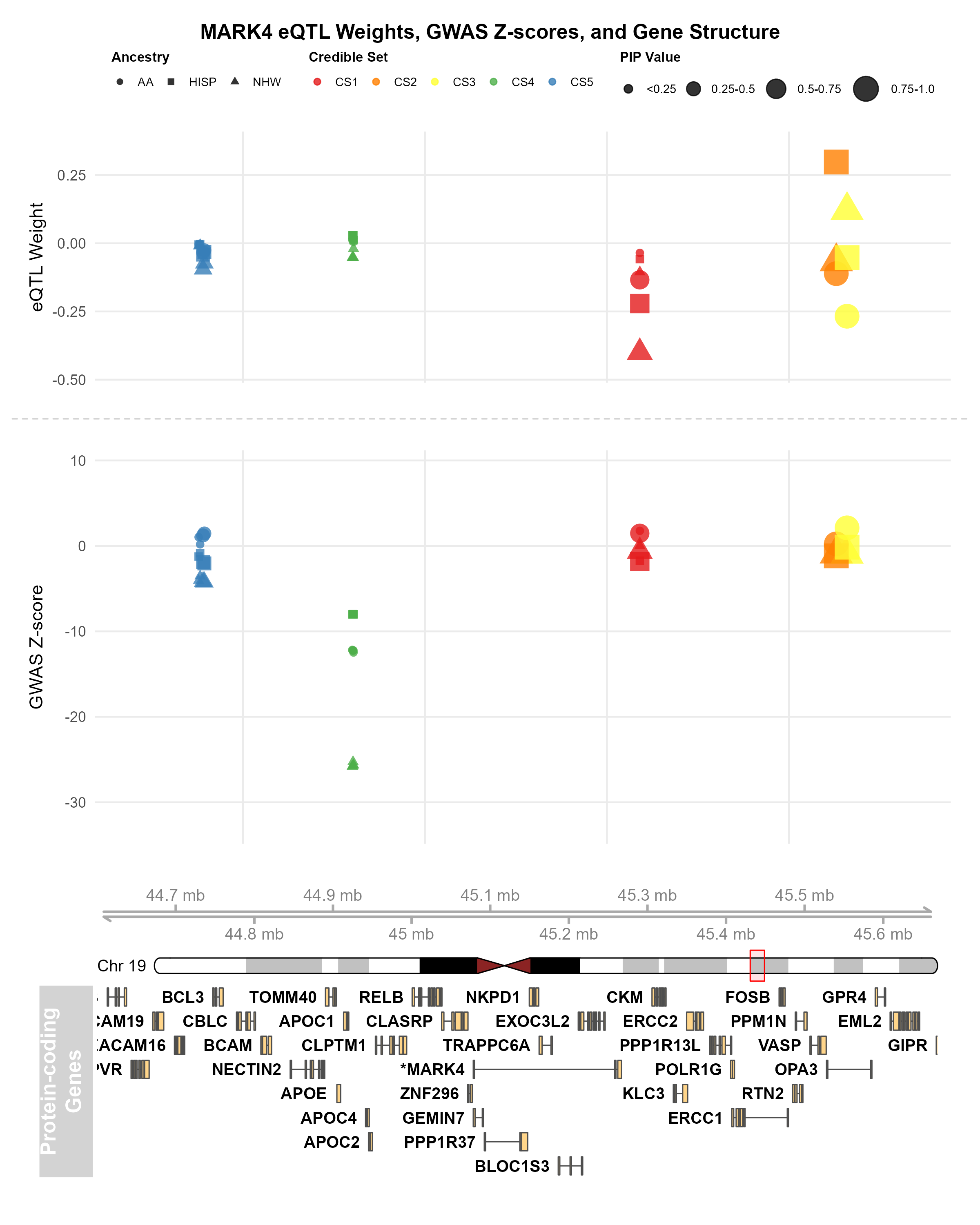

Figure S9: **Fine-mapped eQTLs and GWAS associations for the MARK4 locus.** The figure displays the relationship between eQTL weights (top panel), GWAS Z-scores (middle panel), and gene structure (bottom panel) for MARK4 on chromosome 19. In the top and middle panels, eQTL weights and GWAS associations are shown across three ancestral populations (African American (AA, circles), Hispanic (HISP, squares), and non-Hispanic White (NHW, triangles)), with variants colored by credible set membership. Point sizes indicate posterior inclusion probability (PIP) values. The bottom panel illustrates the genomic structure of MARK4.

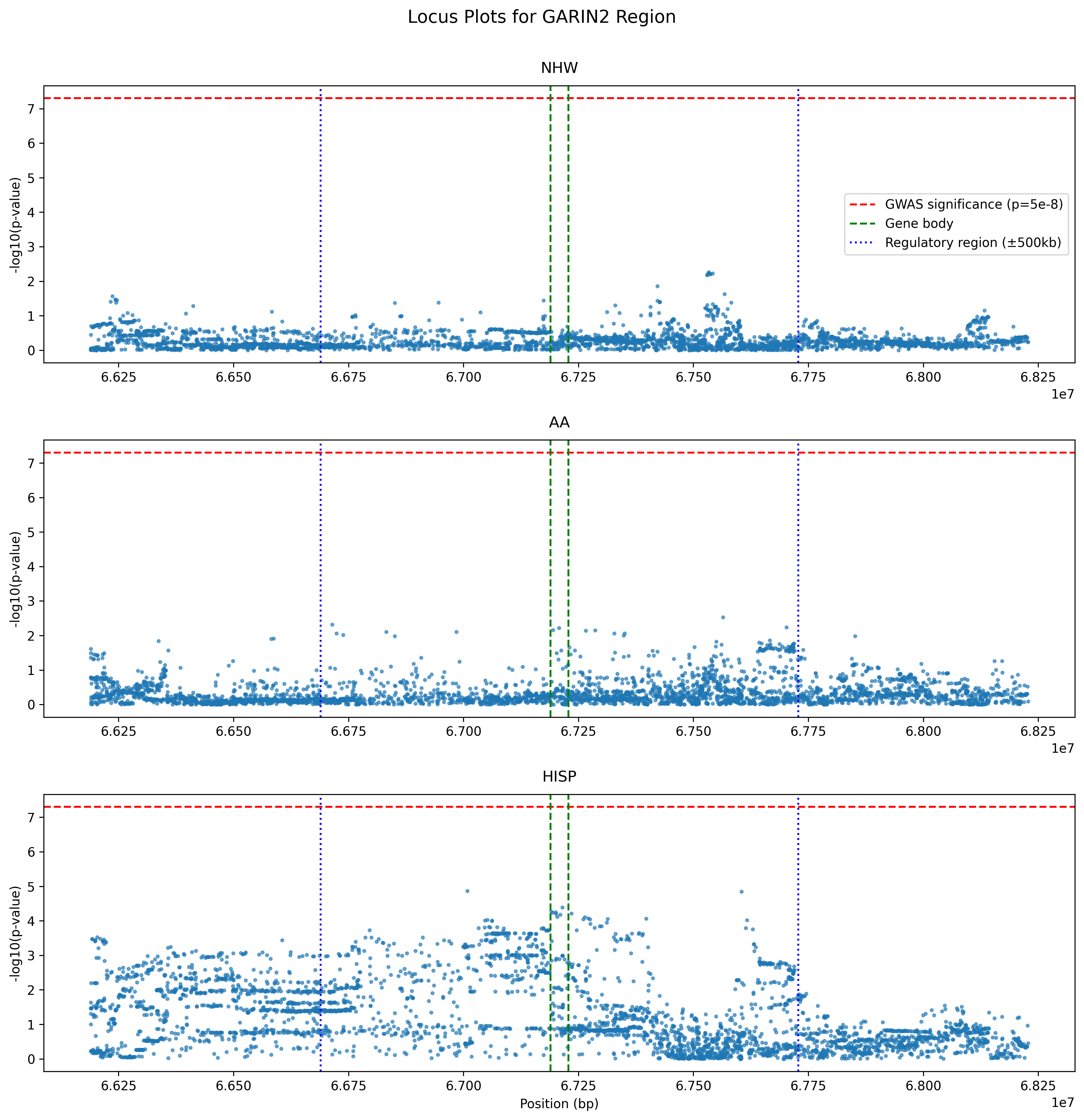

Figure S10: **GWAS associations of cis-regulatory variants near GARIN2 showed suggestive significance in HISP AD GWAS only** The locus plots showed the GWAS associations’ $-{log}_{10}\left( P \right)$ for NHW, AA, and HISP population respectively. HISP showed uniform suggestive significance ($P<1*{10}^{-3}$).

#### Supplemental Tables

**Table S1: Sample characteristics of MAGENTA study participants.**

|  | Subcategory | All | AA | NHW | HISP |
| --- | --- | --- | --- | --- | --- |
| Sample Size (N) |  | 751 | 224 | 235 | 292 |
| Sex | Male | 267 (35.6%) | 64 (28.6%) | 110 (46.8%) | 93 (31.8%) |
|  | Female | 484 (64.4%) | 160 (71.4%) | 125 (53.2%) | 199 (68.2%) |
| Age (years) | Mean ± SD | 77.0 ± 7.7 | 76.6 ± 7.8 | 77.5 ± 7.4 | 76.9 ± 7.9 |
|  | Median [IQR] | 77.0 [71.0-83.0] | 76.5 [70.0-83.0] | 78.0 [72.0-83.0] | 77.0 [71.0-83.0] |
|  | Range | 53.0-98.0 | 63.0-97.0 | 65.0-97.0 | 53.0-98.0 |
| Status | Case | 378 (50.3%) | 111 (49.6%) | 118 (50.2%) | 149 (51.0%) |
|  | Control | 373 (49.7%) | 113 (50.4%) | 117 (49.8%) | 143 (49.0%) |

^a^The table presents demographic and clinical characteristics of the MAGENTA study across all participants and stratified by population groups (AA: African American, NHW: non-Hispanic White, HISP: Hispanic). For each characteristic, counts and percentages are shown for categorical variables, while mean ± standard deviation, median [interquartile range], and range are provided for continuous variables. Sample sizes, sex distribution, age statistics, and Alzheimer’s Disease status are reported for the total cohort and each population group.

Table S2: **Sample characteristics of three genome-wide association studies (GWAS) participants.**

| Population | AD Cases | Controls | Total |
| --- | --- | --- | --- |
| African American | 2,784 | 5,222 | 9,168 |
| Hispanic | 3,005 | 5,894 | 8,899 |
| non-Hispanic White | 21,982 | 44,944 | 63,926 |

^a^African American data from Ray et al.^34^, Hispanic data from Rajabli et al.^4^, and non-Hispanic White data from Kunkle et al.^35^. All summary statistics were derived from models adjusted for age, sex, and population substructure principal components.

Table S3: **SuShiE Fine-Mapping Parameters**

| Parameter | Type | Values |
| --- | --- | --- |
| --L | Integer | 10 |
| --pi | String | “uniform” |
| --resid-var | Float | 1e-3 |
| --effect-var | Float | 1e-3 |
| --rho | Float | 0.1 |
| --no-scale | Boolean | False |
| --no-regress | Boolean | False |
| --no-update | Boolean | False |
| --max-iter | Integer | 500 |
| --min-tol | Float | 1e-3 |
| --threshold | Float | 0.90 |
| --purity | Float | 0.5 |
| --purity_method | String | “weighted” |
| --ld-adjust | Float | 0 |
| --max-select | Integer | 250 |
| --min-snps | Integer | 100 |
| --rint | Boolean | False |
| --no-reorder | Boolean | False |
| --keep-ambiguous | Boolean | False |
| --her | Boolean | True |
| --cv | Boolean | True |
| --cv-num | Integer | 5 |
| --seed | Integer | 12345 |
| --alphas | Boolean | False |

^a^This table provides the complete set of SuShiE fine-mapping parameters employed in our analysis that were not explicitly discussed in the main text. These include algorithm-specific parameters controlling effect estimation, optimization behavior, credible set construction, and cross-validation. All parameters are listed with the values to ensure complete reproducibility of our results.

Table S4: **Population-stratified TWAS results for genes significant in the unadjusted (main) analysis, compared to AD-status-adjusted results (with change in Z-score).**

| Gene Symbol | CHR | Population | TWAS.Z_unadj | TWAS.P_unadj | TWAS.Z_adj | TWAS.P_adj | Z_diff |
| --- | --- | --- | --- | --- | --- | --- | --- |
| *MARK4* | 19 | AA | -4.710 | 2.5e-06 | -2.226 | 2.6e-02 | -2.484 |
| *TOMM40* | 19 | AA | 10.800 | 3.9e-27 | 9.252 | 2.21e-20 | 1.548 |
| *BLOC1S3* | 19 | NHW | 21.200 | 3.46e-100 | 6.960 | 3.41e-12 | 14.240 |
| *TOMM40* | 19 | NHW | 47.700 | 0 | 36.800 | 2.39e-296 | 10.900 |
| *MS4A4E* | 11 | NHW | 7.730 | 1.07e-14 | 5.298 | 1.17e-07 | 2.433 |
| *CEBPZOS* | 2 | NHW | -4.076 | 4.58e-05 | -2.524 | 1.16e-02 | -1.552 |
| *GYPC* | 2 | NHW | -4.433 | 9.28e-06 | -5.537 | 3.08e-08 | 1.103 |
| *CCDC9* | 19 | NHW | -4.140 | 3.47e-05 | -3.390 | 7.03e-04 | -0.750 |
| *DMPK* | 19 | NHW | 5.500 | 3.78e-08 | 4.860 | 1.15e-06 | 0.640 |
| *BIN1* | 2 | NHW | 4.734 | 2.2e-06 | 5.165 | 2.4e-07 | -0.431 |
| *CREBZF* | 11 | NHW | -5.861 | 4.6e-09 | -6.193 | 5.92e-10 | 0.332 |
| *COG4* | 16 | NHW | 4.601 | 4.2e-06 | 4.426 | 9.6e-06 | 0.175 |
| *PTK2B* | 8 | NHW | 4.560 | 5.11e-06 | 4.684 | 2.81e-06 | -0.124 |
| *MYBPC3* | 11 | NHW | 6.283 | 3.33e-10 | 6.297 | 3.04e-10 | -0.014 |
| *TOMM40* | 19 | HISP | -6.752 | 1.46e-11 | -5.536 | 3.1e-08 | -1.217 |
| *GARIN2* | 14 | HISP | -4.622 | 3.8e-06 | -4.521 | 6.14e-06 | -0.101 |

^a^This table summarizes TWAS association statistics for genes that were significant in the main analysis (the model without AD-status adjustment). Results are shown by population (AA, NHW, HISP) and chromosome, reporting unadjusted and AD-status-adjusted TWAS Z-scores and P-values. $Z_{\text{diff}}$ (unadjusted minus adjusted) quantifies attenuation or strengthening of the association after AD adjustment; larger absolute values indicate greater sensitivity to AD-status adjustment, while small differences suggest the signal is robust to this adjustment.

Table S5: **Comparison of SNP inclusion in credible sets between models with and without Alzheimer’s disease adjustment.**

| Credible Set | Total SNPs (AD-adjusted) | Total SNPs (Unadjusted) | Shared SNPs | Shared % (AD-adjusted) | Shared % (Unadjusted) |
| --- | --- | --- | --- | --- | --- |
| CS1 | 88,675 | 89,173 | 67,267 | 75.86% | 75.43% |
| CS2 | 44,556 | 43,302 | 23,583 | 52.93% | 54.46% |
| CS3 | 23,057 | 21,691 | 9,610 | 41.68% | 44.30% |
| Overall | 177,307 | 173,990 | 127,718 | 72.03% | 73.41% |

^a^This table presents the overlap of SNPs identified in credible sets from SuShiE with and without adjustment for Alzheimer’s disease (AD). For each credible set (CS1-CS3, in decreasing order of importance), the table shows the total number of SNPs in each model, the number of shared SNPs between models, and the percentage of SNPs shared. The “Overall” row summarizes statistics across all credible sets. While the highest-ranked credible set (CS1) shows substantial concordance (~75% overlap), the agreement decreases for lower-ranked credible sets.

Table S6: **eQTL effect-size concordance between AD-adjusted and non-adjusted GReX models for all genes and top 5% max|Z| genes.**

| Ancestry | Set | Matched SNP-gene pairs (n) | Weight correlation (r) | Mean \|Δweight\| |
| --- | --- | --- | --- | --- |
| AA | All genes | 27342985 | 0.828 | 0.000127 |
| NHW | All genes | 27346500 | 0.847 | 0.000125 |
| HISP | All genes | 26585527 | 0.869 | 0.000084 |
| AA | Top 5% max\|Z\| genes | 1370321 | 0.784 | 0.000151 |
| NHW | Top 5% max\|Z\| genes | 1310634 | 0.817 | 0.000152 |
| HISP | Top 5% max\|Z\| genes | 1343046 | 0.863 | 0.000098 |

^a^Shared SNP-gene pairs were matched between SuShiE fine-mapping weights files from with and without adjusting AD status for each of the populations. Pearson correlation is computed on matched weights. Mean absolute weight difference is reported as |wunadjusted - wadjusted|.

Table S7: **Genes with large TWAS differences between non-adjusted and AD-adjusted models (|ΔZ| > 4).**

| Ancestry | Gene symbol | CHR | In APOE region | Z without AD adjustment | Z with AD adjustment | \|ΔZ\| | Significant in either model (FDR<0.05) |
| --- | --- | --- | --- | --- | --- | --- | --- |
| AA | SLC39A3 | 19 | No | -1.190 | 2.975 | 4.165 | No |
| NHW | MARK4 | 19 | Yes | 10.900 | 1.010 | 9.890 | Yes |
| NHW | TRAPPC6A | 19 | Yes | 25.100 | 1.950 | 23.150 | Yes |
| NHW | BCL3 | 19 | No | 9.850 | 0.504 | 9.346 | Yes |
| NHW | CLASRP | 19 | Yes | -1.580 | -12.500 | 10.920 | Yes |
| NHW | PPP1R37 | 19 | Yes | 21.500 | 3.860 | 17.640 | Yes |
| NHW | OPA3 | 19 | Yes | 5.020 | 0.521 | 4.499 | Yes |
| NHW | TOMM40 | 19 | Yes | 47.700 | 36.800 | 10.900 | Yes |
| NHW | GEMIN7 | 19 | Yes | -1.090 | -5.380 | 4.290 | Yes |
| NHW | BLOC1S3 | 19 | Yes | 21.200 | 6.960 | 14.240 | Yes |
| HISP | PFKFB2 | 1 | No | 1.398 | -2.694 | 4.092 | No |
| HISP | DOK1 | 2 | No | 2.231 | -1.895 | 4.126 | No |

^a^ΔZ is defined as Zwithout AD adjustment - Zwith AD adjustment. APOE region is defined as chr19:44.8-45.8Mb.

Table S8: **Credible set analysis of 10 AD-related genes analyzed in sparse models**

| Gene | Credible Set | # Variants | Min PIP | Median PIP | Max PIP | Sum PIP |
| --- | --- | --- | --- | --- | --- | --- |
| *BIN1* | 1 | 1 | 1.000 | 1.000 | 1.000 | 1.000 |
| *BIN1* | 2 | 2 | 0.499 | 0.499 | 0.499 | 0.999 |
| *BLOC1S3* | 1 | 10 | 0.018 | 0.124 | 0.124 | 0.917 |
| *COG4* | 1 | 4 | 0.069 | 0.222 | 0.395 | 0.908 |
| *COG4* | 2 | 5 | 0.023 | 0.054 | 0.709 | 0.921 |
| *DMPK* | 1 | 1 | 0.999 | 0.999 | 0.999 | 0.999 |
| *DMPK* | 2 | 8 | 0.086 | 0.090 | 0.185 | 0.941 |
| *DMPK* | 3 | 3 | 0.175 | 0.243 | 0.559 | 0.976 |
| *DMPK* | 4 | 7 | 0.043 | 0.066 | 0.401 | 0.927 |
| *DMPK* | 5 | 52 | 0.007 | 0.020 | 0.037 | 0.901 |
| *DMPK* | 6 | 11 | 0.013 | 0.085 | 0.160 | 0.903 |
| *MARK4* | 1 | 2 | 0.193 | 0.459 | 0.725 | 0.918 |
| *MARK4* | 2 | 1 | 0.979 | 0.979 | 0.979 | 0.979 |
| *MARK4* | 3 | 1 | 0.979 | 0.979 | 0.979 | 0.979 |
| *MARK4* | 4 | 5 | 0.083 | 0.212 | 0.212 | 0.930 |
| *MARK4* | 5 | 4 | 0.034 | 0.212 | 0.472 | 0.930 |
| *MS4A4E* | 1 | 3 | 0.084 | 0.283 | 0.559 | 0.926 |
| *MS4A4E* | 2 | 6 | 0.037 | 0.139 | 0.269 | 0.931 |
| *MYBPC3* | 1 | 34 | 0.008 | 0.022 | 0.116 | 0.907 |
| *PTK2B* | 1 | 6 | 0.059 | 0.171 | 0.238 | 0.903 |
| *PTK2B* | 2 | 9 | 0.078 | 0.089 | 0.166 | 0.932 |
| *PTK2B* | 3 | 12 | 0.012 | 0.036 | 0.474 | 0.904 |
| *TOMM40* | 1 | 1 | 0.998 | 0.998 | 0.998 | 0.998 |
| *TOMM40* | 2 | 2 | 0.382 | 0.497 | 0.613 | 0.994 |
| *TOMM40* | 3 | 1 | 0.950 | 0.950 | 0.950 | 0.950 |
| *TOMM40* | 4 | 9 | 0.005 | 0.034 | 0.352 | 0.903 |
| *TOMM40* | 5 | 15 | 0.011 | 0.023 | 0.410 | 0.911 |

^a^Summary of fine-mapping results for AD-associated genes (complementary to locus plots of 10 significant genes analyzed in sparse models). The Min/Median/Max PIP columns show the distribution of posterior inclusion probabilities within each set, while Sum PIP indicates the cumulative PIP captured by the credible set.

Table S9: **Cross-population cis-heritability and predictive performance of expression models**

| Gene | h²_AA | R²_AA | h²_NHW | R²_NHW | h²_HISP | R²_HISP |
| --- | --- | --- | --- | --- | --- | --- |
| *BIN1* | 0.0312* | 0.1787*** | 0.0562*** | 0.2626*** | 0.0338* | 0.1165*** |
| *TOMM40* | 0.0000 | -0.0015 | 0.0448** | 0.1326*** | 0.0408 | -0.0010 |
| *DMPK* | 0.0824* | 0.1459*** | 0.0158 | 0.0683*** | 0.0274 | 0.0259** |
| *PTK2B* | 0.0200 | 0.0576*** | 0.0154 | 0.1197*** | 0.0284 | 0.0399*** |
| *MYBPC3* | 0.0080 | 0.0181* | 0.0520*** | 0.1683*** | 0.0250 | 0.0013 |
| *CREBZF* | 0.0020 | -0.0043 | 0.0004 | -0.0025 | 0.0111 | 0.0013 |
| *BLOC1S3* | 0.0000 | -0.0041 | 0.0483** | 0.0312** | 0.0164 | 0.0055 |
| *MS4A4E* | 0.0173 | 0.0501*** | 0.0002 | 0.0004 | 0.0000 | -0.0032 |
| *COG4* | 0.0646*** | 0.1227*** | 0.0193* | 0.0487*** | 0.0000 | 0.0112* |
| *MARK4* | 0.0000 | 0.0033 | 0.1373*** | 0.3079*** | 0.0358 | 0.1017*** |

^a^Per-gene cis-SNP heritability (h²) and 5-fold cross-validated prediction R² (R²) reported by population (AA, NHW, HISP). Asterisks denote nominal two-sided significance for the corresponding entry (*** $P<0.001$, ** $P<0.01$, * $P<0.05$; no symbol = not significant). Values are shown to four decimals; significance is evaluated on unrounded P-values.

Table S10: **MA-FOCUS PIP of significant TWAS associations**

| gene_symbol | PIP_NHW | PIP_AA | PIP_HISP | PIP_ME |
| --- | --- | --- | --- | --- |
| *TOMM40* | 1.000 | 1.000 | 1.000 | 1.000 |
| *BIN1* | 1.000 | 0.857 | 0.140 | 1.000 |
| *BLOC1S3* | 1.000 | 0.006 | 0.029 | 1.000 |
| *MS4A4E* | 0.995 | 0.002 | 0.001 | 0.998 |
| *MYBPC3* | 0.990 | 0.004 | 0.008 | 0.996 |
| *CREBZF* | 1.000 | 0.011 | 0.017 | 0.972 |
| *DMPK* | 0.863 | 0.003 | 0.004 | 0.916 |
| *GARIN2* | 0.030 | 0.039 | 0.908 | 0.883 |
| *PTK2B* | 0.978 | 0.027 | 0.025 | 0.866 |
| *CEBPZOS* | 0.928 | 0.011 | 0.013 | 0.596 |
| *COG4* | 0.969 | 0.003 | 0.004 | 0.418 |
| *CCDC9* | 0.855 | 0.003 | 0.003 | 0.245 |
| *MARK4* | 0.130 | 0.955 | 0.005 | 0.020 |

^a^Dense-model TWAS genes significant by both MA-FOCUS (PIP $\geq$ 0.8) and FUSION TWAS ($q_{c}<0.05$), showing the maximum PIP for NHW, AA, HISP, and cross-population (ME).

Table S11: **Validation of TWAS associations using eQTL reference panels from TOPMed MESA visit 1 by population**

| Gene | Z AA | Z NHW | Z HISP |
| --- | --- | --- | --- |
| *BIN1* | 2.18* | 2.62** | 0.99 |
| *TOMM40* | 9.57*** | 15.30*** | -3.64*** |
| *DMPK* | 0.28 | 4.28*** | 2.96** |
| *PTK2B* | 0.00 | 4.49*** | 0.38 |
| *MYBPC3* | -0.06 | 5.78*** | 0.60 |
| *CREBZF* | 1.20 | -1.04 | -0.92 |
| *BLOC1S3* | -0.17 | -4.43*** | 0.01 |
| *MS4A4E* | 0.91 | 6.72*** | -1.97* |
| *COG4* | -0.18 | 0.28 | 0.76 |
| *MARK4* | -1.66 | 0.51 | -1.17 |

^a^Per-gene TWAS Z-scores from the TOPMed MESA visit 1 validation dataset, stratified by population (AA, NHW, HISP). Asterisks denote nominal two-sided significance for the corresponding entry (*** $P<0.001$, ** $P<0.01$, * $P<0.05$; no symbol = not significant).

Table S12: **Cross-validated predictive performance (R²) of dense and sparse eQTL prediction models across three ancestral populations**

| Ancestry | n (genes) | Dense R² Mean (SE) | Dense R² Median | Sparse R² Mean (SE) | Sparse R² Median |
| --- | --- | --- | --- | --- | --- |
| AA | 8887 | 0.0300 (0.0005) | 0.0114 | 0.0346 (0.0005) | 0.0167 |
| NHW | 8921 | 0.0525 (0.0009) | 0.0174 | 0.0524 (0.0008) | 0.0225 |
| HISP | 8698 | 0.0238 (0.0007) | 0.0036 | 0.0235 (0.0006) | 0.0050 |

^a^Comparison of out-of-sample R² between dense models using all cis-SNPs and sparse models using only fine-mapped SNPs in credible sets. The dense model includes all cis-SNPs within the cis-region, while the sparse model uses only variants identified through fine-mapping (credible set SNPs from the main analysis). Results are shown for African American (AA; n = 8,887 genes), non-Hispanic White (NHW; n = 8,921 genes), and Hispanic (HISP; n = 8,698 genes) populations. Values represent mean R² with standard error (SE) in parentheses, and median R². Across all populations, sparse models achieved comparable or higher predictive performance than dense models, demonstrating that fine-mapped causal eQTL variants effectively capture the predictive signal without requiring all cis-SNPs.

Table S13: **COG4 credible set variant frequencies across ancestries**

| SNP ID | CS | NHW Freq | AA Freq | HISP Freq |
| --- | --- | --- | --- | --- |
| chr16:70508818_C_T | 1 | 0.034 | 0.241 | 0.036 |
| chr16:70616971_G_A | 1 | 0.038 | 0.228 | 0.045 |
| chr16:70617075_T_C | 1 | 0.038 | 0.228 | 0.045 |
| chr16:70470982_T_C | 1 | 0.034 | 0.248 | 0.045 |
| chr16:70666311_A_C | 2 | 0.496 | 0.174 | 0.380 |
| chr16:70698580_G_A | 2 | 0.521 | 0.143 | 0.445 |
| chr16:70696670_A_C | 2 | 0.521 | 0.143 | 0.447 |
| chr16:70690022_A_G | 2 | 0.575 | 0.413 | 0.505 |
| chr16:70669915_T_G | 2 | 0.466 | 0.179 | 0.435 |

^a^SNP ID uses chr{chr}:pos_ref_alt. CS is the SuShiE credible set index. NHW/AA/HISP Freq are alternative allele frequencies from each group.

Table S14: **Functional annotation of COG4 eQTL CS2 variants**

| Variant | PIP | ChromHMM State(s) | No. Cell Types | pcHi-C to *COG4* |
| --- | --- | --- | --- | --- |
| **chr16:70666311_A_C** | 0.709 | EnhA2 | 1 (GM19238) | Detected (2.65)^a^ |
| chr16:70698580_G_A | 0.111 | EnhA1/A2/G1/G2/Wk | 18 | - |
| chr16:70696670_A_C | 0.054 | EnhA1/Wk | 34 | - |
| chr16:70690022_A_G | 0.024 | EnhA1/A2/Wk | 16 | - |
| chr16:70669915_T_G | 0.023 | - | 0 | Detected (2.65)^a^ |

^a^Variants in the second credible set for *COG4* expression were annotated using chromatin state predictions from EpiMap ChromHMM and promoter-capture Hi-C (pcHi-C) interaction data from Javierre et al. (2016). Posterior inclusion probabilities (PIPs) were derived from multi-ancestry SuSiE fine-mapping. ChromHMM enhancer states include active enhancers (EnhA1, EnhA2), genic enhancers (EnhG1, EnhG2), and weak enhancers (EnhWk) across blood and immune cell types. pcHi-C interactions represent physical chromatin contacts between the variant-containing region (chr16:70652543-70673594) and the *COG4* promoter; CHiCAGO scores $<$ 5 indicate detection below the statistical significance threshold.

Table S15: **CHiCAGO scores for pcHi-C contacts between the lead-variant region and the COG4 promoter**

| Cell Type | CHiCAGO Score |
| --- | --- |
| Macrophage (Mac0) | 2.65 |
| CD8+ naive T cell | 2.45 |
| Monocyte | 2.39 |
| CD8+ T cell (total) | 1.38 |
| CD4+ naive | 1.32 |
| CD4+ (total) | 1.05 |
| Naive B cell | 0.37 |
| Neutrophil | 0.16 |

^a^Promoter-capture Hi-C (pcHi-C) in primary hematopoietic cells shows detectable but sub-threshold chromatin contact between the lead-variant region (chr16:70,652,543-70,673,594) and the *COG4* promoter (chr16:70,522,638-70,524,465). The scores do not meet the significance threshold (CHiCAGO ≥ 5).
